# T Cell Deficiency Precipitates Antibody Evasion and Emergence of Neurovirulent Polyomavirus

**DOI:** 10.1101/2022.02.25.481945

**Authors:** Matthew D. Lauver, Ge Jin, Katelyn N. Ayers, Sarah N. Carey, Charles S. Specht, Catherine S. Abendroth, Aron E. Lukacher

## Abstract

JC polyomavirus (JCPyV) causes Progressive Multifocal Leukoencephalopathy (PML), a life-threatening brain disease in T cell immunosuppressed patients. PML patients often carry mutations in the JCPyV VP1 capsid protein. These mutations confer resistance to neutralizing VP1 antibodies (Ab). We found that T cell insufficiency during persistent infection, in the setting of monospecific VP1 Ab, was required for outgrowth of VP1 Ab-escape viral variants. CD4 T cells were primarily responsible for preventing resurgent virus infection in the kidney and checking emergence of these mutant viruses. T cells also provided a second line of defense against Ab-escape VP1 mutant viruses. A virus with two capsid mutations, one conferring Ab-escape yet impaired infectivity and a second compensatory mutation, yielded a highly neurovirulent variant. These findings link T cell deficiency and evolution of Ab-escape polyomavirus VP1 variants with neuropathogenicity.

## Introduction

Antibodies are critical components of immune defense against viral pathogens and key mediators of immune control during persistent infections. Evasion of antibodies, particularly in settings of limited antibody diversity, is a potent selective pressure for viral mutations. Notably, outgrowth of escape variants is an Achilles’ heel for monoclonal antibody antiviral intervention (Bar et al., 2016; Caskey et al., 2015, 2017; Mehandru et al., 2007; Toma et al., 2011; Trkola et al., 2005).

Intrahost viral evolution to select variants that evade neutralizing antibody is common among chronic viral infections. RNA viruses typically exist as a quasispecies generated by their error-prone viral polyomerases, a viral lifestyle that provides a cocktail of variants to probe for resistance against antiviral antibodies (Inuzuka et al., 2018; Kinchen et al., 2018; Lynch et al., 2015). DNA viruses, however, are less prone to existing as a quasispecies, especially those that commandeer the high-fidelity host DNA replication machinery. Evolution by DNA viruses, and sculpting by antiviral antibodies, may be a feature shared by viruses that persistent in their hosts in a “smoldering” infectious state.

Polyomaviruses (PyV) are nonenveloped, dsDNA viruses that infect the majority of the animal kingdom. Highly species-specific, most polyomaviruses cause lifelong, silent infections in their natural hosts. Fourteen human polyomaviruses have been identified, four of which cause severe disease in immunocompromised individuals. Among these is JCPyV, which typically persists asymptomatically in the kidneys and urinary tract, but is also the causative agent of Progressive Multifocal Leukoencephalopathy (PML). PML, a frequently fatal demyelinating brain disease, is associated with various forms of T cell immunosuppression including HIV/AIDS, organ transplant immunosuppressants, and certain immunomodulatory therapies (Cortese et al., 2021; Pavlovic et al., 2018). Typically identified by MRI following the onset of neurologic symptoms, PML is diagnosed at a point when limited treatment options exist and permanent CNS damage has often already occurred. Defining the early stages of disease involved in the transition of JCPyV from a kidney to brain pathogen would thus facilitate earlier detection and therapeutic intervention before the development of neurologic disease.

Although PML progresses rapidly following the emergence of neurological disease, prolonged immune suppression is a necessary antecedent to PML development. For instance, high risk for natalizumab-associated PML involves infusion therapy for >24 months and a history of immunosuppression (Berger and Fox, 2016; Bloomgren et al., 2012; Fox and Rudick, 2012). This long period of immunosuppression before clinical disease manifests raises the possibility that variants of JCPyV, including those with neurovirulent potential, may emerge over time. Supporting this possibility is evidence that JCPyV exists as a quasispecies in the blood and CSF, but not urine, of PML patients (Van Loy et al., 2015). Iatrogenic T cell and B cell ablation therapies have independently been associated with PML. Lacking is a conceptual connection between T cell deficiency, B cell deficiency, and prolonged immunosuppression in JCPyV evolution that could lead to outgrowth of neurovirulent JCPyV variants.

PML is characterized by emergence of mutations in the major capsid protein, VP1, of JCPyV, which are not found in the circulating (i.e., archetype) strains (Gorelik et al., 2011; Zheng et al., 2005b, 2005a). The nonenveloped JCPyV icosahedral capsid is comprised of 360 copies of VP1 organized into 72 pentamers. VP1 mediates attachment to cellular receptors via its four solvent-exposed loops; these loops are also the dominant targets of the host’s neutralizing antibody response (Buch et al., 2015; Lindner et al., 2019; Neu et al., 2010). JCPyV-PML VP1 mutations, which are situated in these loops, mediate resistance to neutralizing antibodies, but also alter receptor binding and viral tropism (Geoghegan et al., 2017; Gorelik et al., 2011; Jelcic et al., 2015; Lauver et al., 2020; Maginnis et al., 2013; Ray et al., 2015). Sera from PML patients as well as some healthy individuals contain “blind spots” for particular JCPyV-PML VP1 mutations, despite these individuals having high antibody titers against WT JCPyV (Ray et al., 2015). The conditions that lead to the outgrowth of these variants in PML patients remain poorly understood, in part due to the lack of animal models for studying the early stages of JCPyV pathogenesis.

We previously identified a neutralizing monoclonal antibody (mAb) against mouse polyomavirus (MuPyV) VP1 (Swimm et al., 2010). The epitope for this mAb is the dominant target of the endogenous VP1 Ab response in mice. Importantly, this mAb failed to recognize several MuPyV VP1 mutations that correspond to mutations seen in JCPyV isolates from PML patients (Lauver et al., 2020). In this study, we utilized passive immunization with this mAb in Ab-deficient mice to mimic the serologic susceptibility of PML patients to JCPyV-PML VP1 variant viruses. We found that T cell loss during persistent infection in these mice led to outgrowth of viruses carrying mutations in similar regions of VP1 to those seen in PML patients. Although an array of VP1 mutations arose during resurgent replication in the kidneys in T cell-depleted hosts, evading Ab neutralization was the primary driver selecting replication-competent mutant viruses, including rare neuropathogenic variants. Thus, T cell deficiency coupled with limited Ab coverage of VP1 epitopes are required for outgrowth of Ab-escape viral variants, including those with a potential for neurovirulence.

## Results

### T cell deficiency during persistent infection leads to outgrowth of antibody-resistant virus variants

We demonstrated that Ab-escape VP1 mutations emerge after serial passage of wild type (WT) MuPyV (strain A2) in host cells in the presence of a VP1-specific neutralizing mAb (Lauver et al., 2020). To ask whether VP1 antibody-escape mutations similarly arise in vivo, we passively immunized B cell-deficient μMT mice with this VP1-specific mAb (clone 8A7H5) to model serologic “blindness” to particular JCPyV VP1 mutations seen in PML patients (Ray et al., 2015). MuPyV-infected μMT mice developed high-titer chronic viremia, which was efficiently controlled by injection of 8A7H5 mAb starting at 4 days post-infection (dpi) and continuing weekly thereafter (**Figure S1A-B**). At 20 dpi, sera of VP1 mAb-treated μMT mice displayed similar neutralizing titers as B6 mice against WT MuPyV, but an inability to neutralize a VP1 mutant virus with a deletion of aspartic acid at position 295, A2.Δ295, which we previously identified as being resistant to 8A7H5 (Lauver et al., 2020)(**Figure S1C-E**).

We first investigated the effect of T cell deficiency on virus control. CD4 and/or CD8β T cell-depleting antibodies or control rat IgG were administered to VP1 mAb-treated μMT mice beginning at 20 dpi (**Figure 1A**). Ten days after starting T cell depletion, virus levels were significantly elevated in the kidney, the dominant site of MuPyV persistent infection, in mice receiving combined T cell depletion (**Figure 1B**). In mice receiving CD4 or CD8β T cell-depleting antibodies individually, kidney virus levels were increased in CD4, but not CD8β, T cell depleted mice (**Figure 1C**). In dual CD4 and CD8 T cell depleted mice, anti-VP1 immunofluorescence staining was localized to disrupted tubules expressing Tamm-Horsfall protein (THP), identifying the distal convoluted tubules as the major site of MuPyV replication in the kidney (Tokonami et al., 2018)(**Figure 1D**). These findings indicate that T cells prevent resurgence of persistent MuPyV infection from kidney reservoirs.

**Figure 1.**
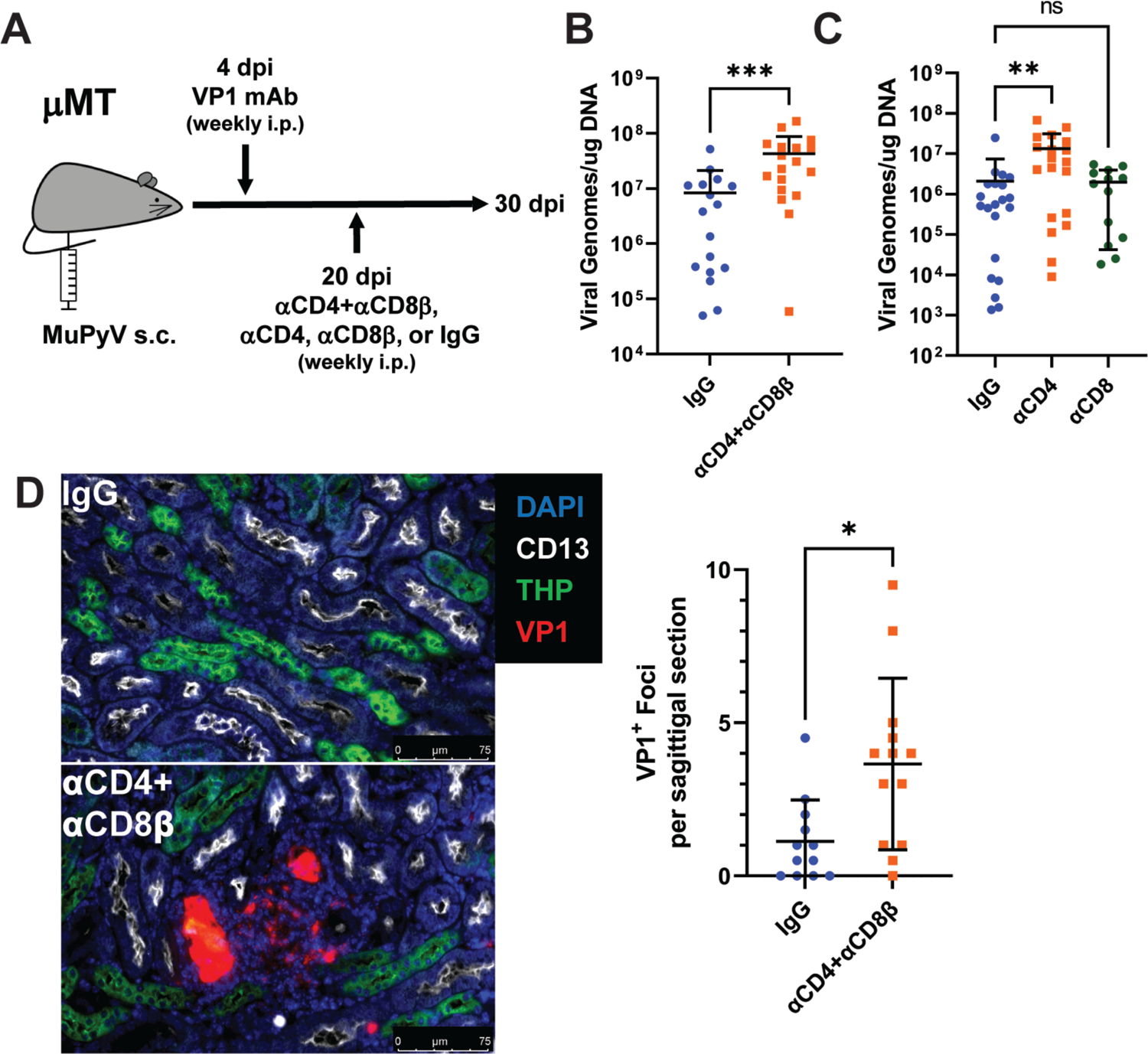
MuPyV resurgence in the kidney following T cell depletion in VP1 mAb-treated μMT mice. A. Experimental approach for VP1 mAb treatment and T cell depletion. B. Viral DNA levels in organs 10 days post T cell depletion with combined αCD4 and αCD8β. Viral DNA was quantified by qPCR and compared to a standard curve (n=16-18). C. Viral DNA levels in organs 10 days post depletion with αCD4 or αCD8β. Viral DNA was quantified by qPCR and compared to a standard curve (n=13-21). D. (Left) Foci of virus replication the kidney cortex 10 days post T cell depletion. Kidneys were stained for CD13 (white), THP (green) and VP1 (red). (Right) Quantification of virus foci in the kidney. Data are the average of two kidney sections per mouse (n=12-13). Error bars are mean ± SD. Data are from at least two independent experiments. Data were analyzed by Mann-Whitney U test (B, D) or Kruskal-Wallis test with Dunn’s multiple comparisons test (C). *p < 0.05, **p < 0.01, ***p < 0.001. See also **Figure S1**.

Next, we asked how T cell loss affected long term virus control. Mice were treated with combined CD4 and CD8β T cell-depleting antibodies, CD4 T cell-depleting antibody, CD8β T cell-depleting antibody, or control rat IgG. Blood was collected every 20 days and screened for infectious (i.e., un-neutralized) virus by plaque assay (**Figure 2A**). Viremia became detectable in CD4 and CD8β T cell-depleted mice as early as 40 days after dual CD4 and CD8 T cell depletion, with all mice developing viremia by approximately 80 days post depletion. CD4 T cell loss alone also led to viremia development in all mice by 120 days post depletion (**Figure 2B**). In contrast, fewer than half of persistently infected mice given control IgG or CD8β T cell-depleting antibody developed viremia; in IgG-treated mice the development of viremia was delayed and peaked at approximately 100-fold lower levels than the T cell-depleted mice (**Figure 2B**). Viremic T cell-depleted mice showed high systemic viral infection in the kidney, spleen, and brain (**Figure 2C-E**).

**Figure 2.**
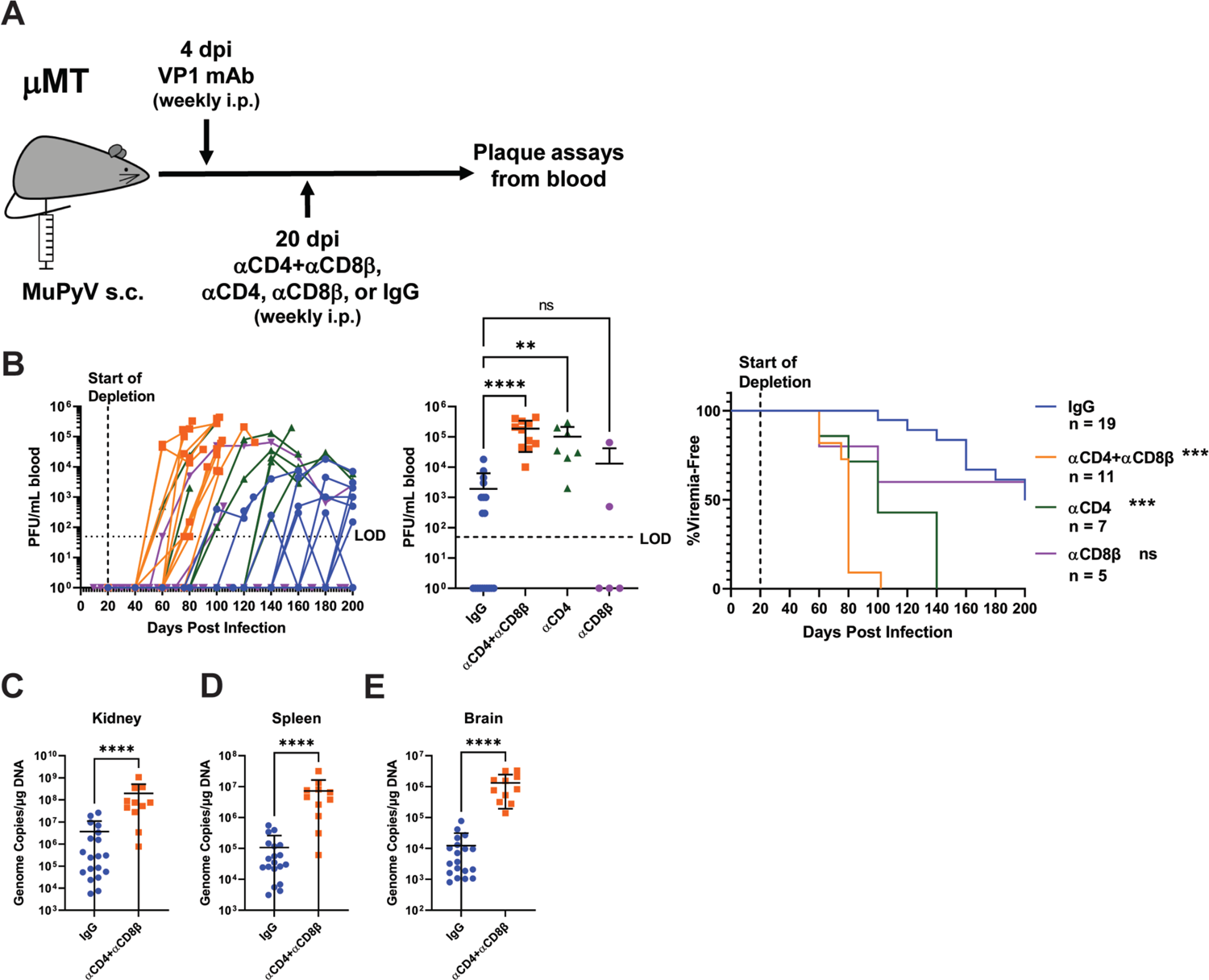
T cell loss enables viremia. A. Experimental scheme for VP1 mAb treatment, T cell depletion, and detection of viremia. B. (Left) Viral titers in the blood of control and T cell depleted mice over time. Viremia was measured by plaque assay from whole blood. (Center) Peak levels of viremia detected in control and T cell depleted mice. (Right) Time to development of viremia in control and T cell depleted mice. Indicated significances are with comparison to the IgG group. LOD: Limit of detection (n=5-19). C-E. Viral DNA levels in the kidney (C), spleen (D), and brain (E) at the time of euthanasia in control and T cell depleted mice. Viral DNA was quantified by qPCR and compared to a standard curve (n=11-19). Error bars are mean ± SD. Data are from at least two independent experiments. Data were analyzed by Kruskal-Wallis test with Dunn’s multiple comparisons test (B Center), Mantel-Cox test with Bonferroni’s correction for multiple comparisons (B Right), Mann-Whitney U test (C-E).**p < 0.01, ***p < 0.001, ****p < 0.0001.

Sequencing plaque-purified virus from the blood of each viremic mouse revealed mutations in VP1, with only one VP1 variant found in individual mice (**Figure 3A** and **Table S1**). This finding mirrors evidence that a PML patient typically harbors a single VP1 mutant virus in their CSF, brain, and blood (Gorelik et al., 2011; Reid et al., 2011; Zheng et al., 2005a). Several of these non-synonomous single nucleotide substitutions and codon deletions were previously isolated from serially passaged virus refractory to 8A7H5 mAb-mediated neutralization, including Δ295, the most frequently detected mutation, as well as V296F, previously identified as the MuPyV equivalent of the common JCPyV PML mutation S268F (Lauver et al., 2020; Sunyaev et al., 2009). Notably, the codon for phenylalanine in V296F in this host-derived MuPyV differs from the one we previously created by site directed mutagenesis (Lauver et al., 2020). We also identified several new mutations, including combined single amino acid deletions and substitutions. Each of the VP1 mutant viruses contained a mutation in the HI loop, where the heavy chain of the 8A7H5 mAb contacts multiple residues (**Figure 3B**) (Lauver et al., 2020). We introduced several of these single and dual VP1 mutations into WT MuPyV and found that they blocked neutralization by 8A7H5 (**Figure 3C**). These mutations, however, had varying effects on the efficiency of viral infection in the kidney and spleen (**Figure 3D-E**). These data indicate that escape from neutralizing antibody, not shifts in tissue tropism, was the selective pressure behind the emergence of VP1 mutations *in vivo*.

**Figure 3.**
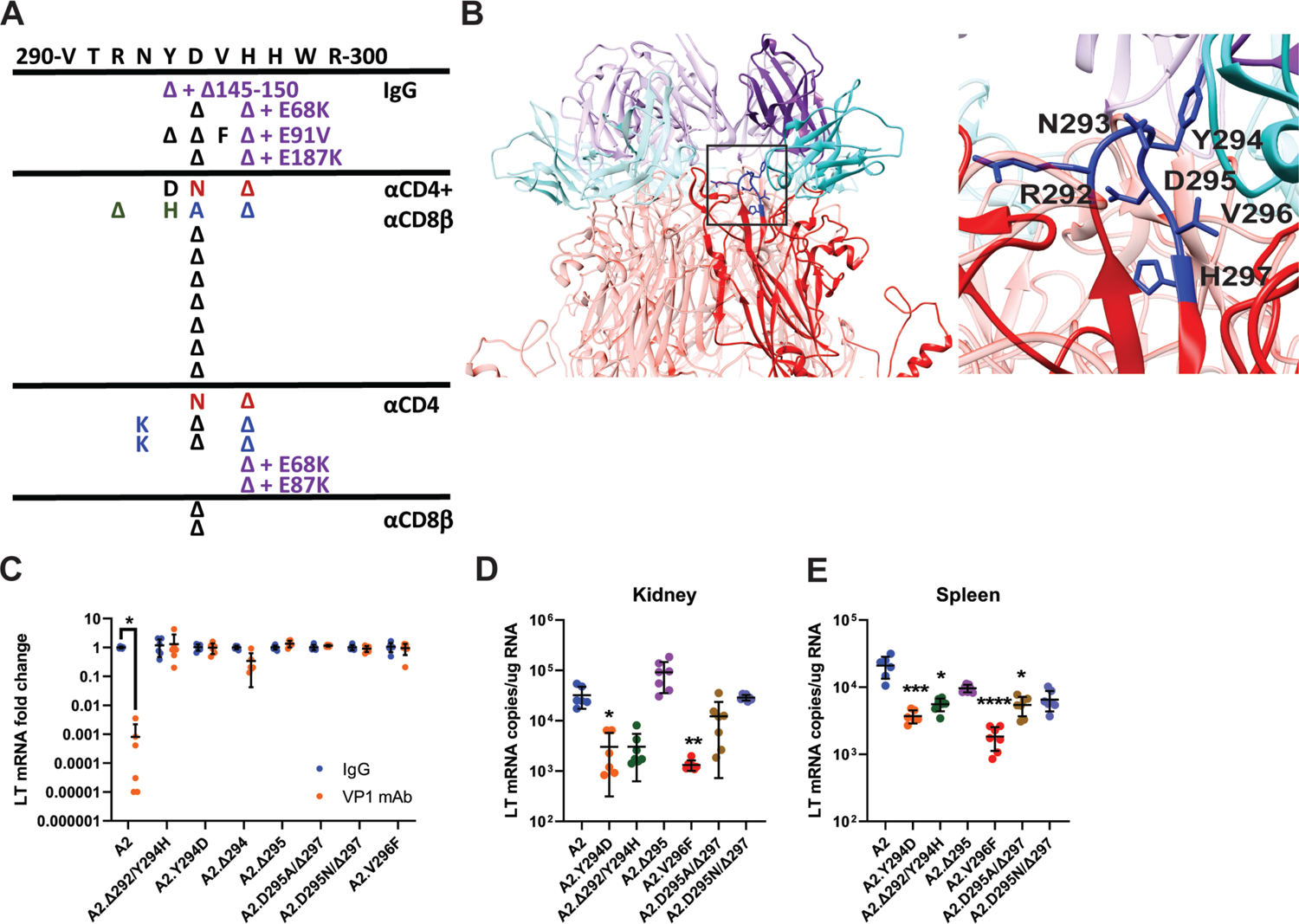
Viremia is mediated by VP1 Ab escape mutations. A. Location of detected VP1 mutations in the HI loop ordered by treatment group. Amino acid sequence from V290 to R300 is listed at the top. Black letters indicate a single mutation, colored letters indicate pairs of mutations in the same virus. Purple lettering designates viruses with a second mutation in a separate loop of VP1. Amino acid deletions are indicated with a Δ. B. Location of VP1 mutations (blue) in the HI loop of one copy of VP1 relative to the location of VP1 mAb (cyan/purple) in the Cryo-EM structure of WT VP1 and the VP1 mAb. PDB ID: 7K22 (Lauver et al., 2020). C. VP1 mAb neutralization of VP1 mutant viruses. Viruses were preincubated with 10μg VP1 mAb or control IgG for 30 min prior to addition to 1 x 10^5^ NMuMG epithelial cells. A2 was diluted to an MOI of 0.1 PFU/cell, mutant viruses were diluted to match A2 by genomic equivalents (g.e.). Viral LT mRNA levels were quantified 24 hpi and normalized for each virus to infection with control IgG (n=6). D-E. Viral mRNA levels in the kidney (E) and spleen (F) 4 dpi with VP1 mutant viruses compared to WT. WT mice were infected i.v. with 1 x 10^6^ PFU of A2 or mutant viruses matched by g.e. Viral LT mRNA levels were quantified by qPCR and compared to a standard curve (n=6-7). Error bars are mean ± SD. Data are from at least two independent experiments. Data were analyzed by Mann-Whitney U test with Holm-Šídák correction for multiple comparisons (C) and Kruskal-Wallis test with Dunn’s multiple comparisons test (D-E).*p < 0.05, **p < 0.01, ***p < 0.001, ****p < 0.0001. See also **Table S1**.

### T cells prevent outgrowth of antibody-escape mutant virus

The absent/delayed and lower viremia in the IgG-treated mice than T cell-depleted mice led us to hypothesize that T cells could prevent/restrain the outgrowth of an antibody-escape virus if one arose during persistent infection. To test this possibility, we treated μMT with VP1 mAb as before, but challenged the mice with 10^3^ PFU of the A2.Δ295 mutant virus two days after starting T cell depletion (**Figure 4A**). Mice receiving CD4 and CD8β T cell-depleting antibodies developed viremia with progressively increasing infectious virus titers over time; in contrast, no viremia was detected in the control IgG-treated mice (**Figure 4B**). Moreover, the T cell-depleted mice had 100-1,000-fold higher virus levels in the kidney, spleen, and brain than the IgG-treated mice (**Figure 4C-E**). These results provide clear evidence that T cells act to prevent viremia by MuPyV variants that escape neutralizing antibody.

**Figure 4.**
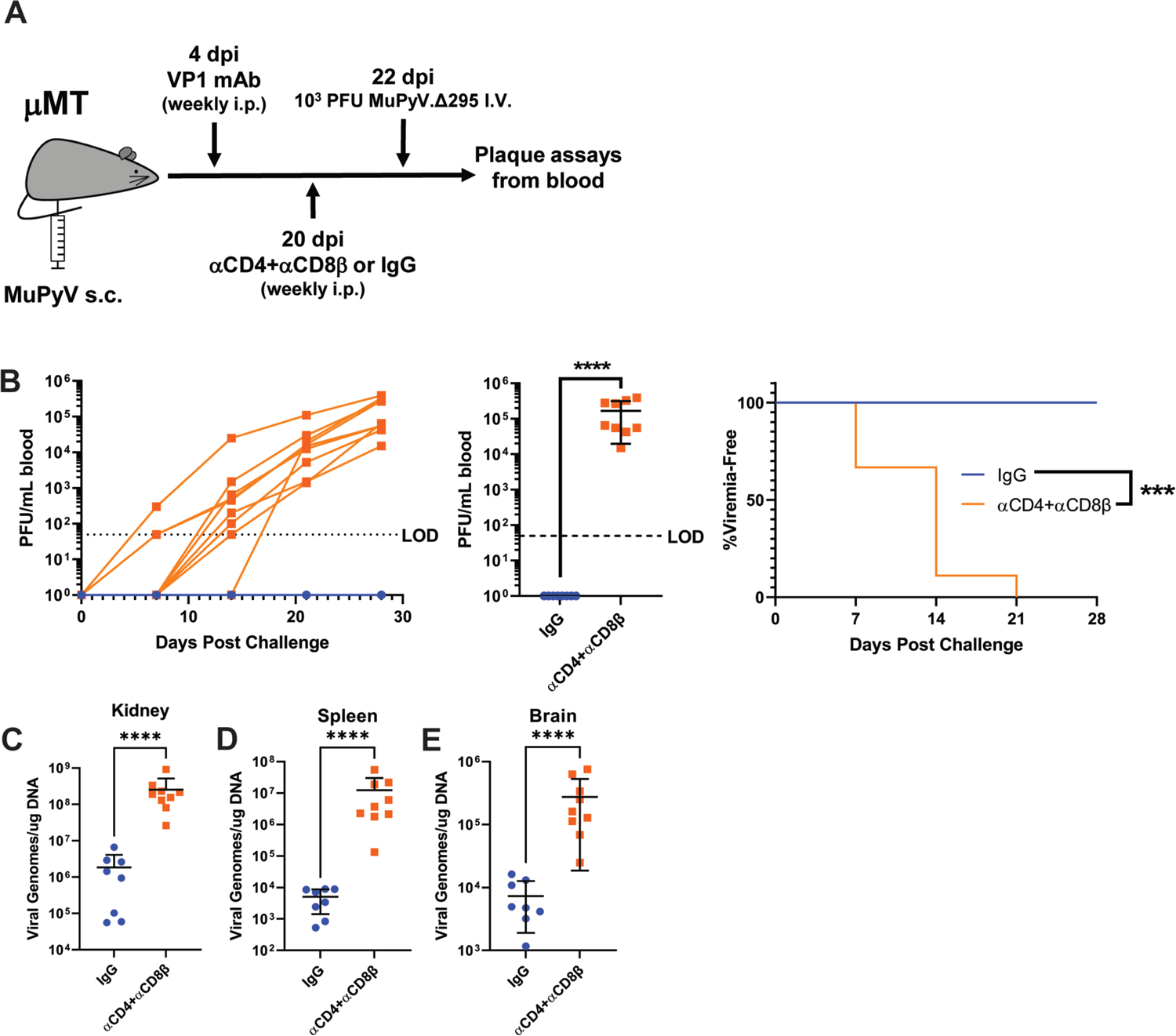
T cells prevent the outgrowth of Ab escape mutant virus. A. Experimental scheme for VP1 mAb treatment, T cell depletion, A2.Δ295 challenge, and detection of viremia. B. (Left) Viral titers in the blood of control and T cell depleted mice over time. Viremia was measured by plaque assay from whole blood. (Center) Peak levels of viremia detected in control and T cell depleted mice. (Right) Time to development of viremia in control and T cell depleted mice in B. LOD: Limit of detection (n=8-9). C-E. Viral DNA levels in the kidney (C), spleen (D), and brain (E) 28 days post challenge (n=8-9). Error bars are mean ± SD. Data are from two independent experiments. Data were analyzed by Mantel-Cox test (B Right) and Mann-Whitney U test (B center, C-E). ***p < 0.001, ****p < 0.0001.

### A double VP1 mutation balances viral fitness vs antibody-escape

Antibody-escape viruses carrying a substitution at D295 together with a deletion of H297 (A2.D295N/Δ297) stand out because the D295 and H297 side chains face away from the interface of VP1 and the 8A7H5 mAb (**Figure 3B**). We generated viruses individually carrying the D295N (A2.D295N) or Δ297 (A2.Δ297) mutations to define the contributions of each mutation to recognition by 8A7H5 and potential effects on tropism. Neutralization assays with 8A7H5 showed that A2.D295N remained sensitive, but A2.Δ297 was fully resistant, to neutralization (**Figure 5A**). Similar to parental A2, A2.D295N was readily bound by 8A7H5; in contrast, 8A7H5 did not bind A2.Δ297 or A2.D295N/Δ297 (**Figure 5B**). A2.D295N showed a similar avidity profile to parental A2 (**Figure 5C**). Taken together, these data show that D295N did not affect 8A7H5 neutralization, suggesting that this mutation did not contribute to antiviral antibody escape. In support, we found that A2.Δ297 failed to form plaques, indicating a defect in spread (**Figure S2A**). In contrast, A2.D295N and A2.D295N/Δ297 formed plaques comparably to WT virus. By titering infectious virus output during one round of replication, we observed decreased virion production by A2.D295N but significantly increased replication by A2.D295N/Δ297 (**Figure 5D**). Matching virus titers by DNA genome equivalents (g.e.), we found that by frequency of early expression of Large T antigen by A2.D295N/Δ297 significantly exceeded that of A2, while A2.Δ297 was similar to A2 and A2.D295N was significantly lower (**Figure 5E**). After a low MOI infection, A2.Δ297 and A2.D295N showed a significant reduction in spread compared to A2 (**Figure 5F**). This reduction in spread not was due to reduced virus production, cells transfected with equal amounts of viral DNA showed similar levels of virus output at 72 h by A2.D295N and A2.Δ297, but significantly more virus by A2.D295N/Δ297 (**Figure S2B**). These data indicate the Δ297 mutation impaired viral spread in a manner independent of virus production, which was rescued by the presence of the D295N mutation.

**Figure 5.**
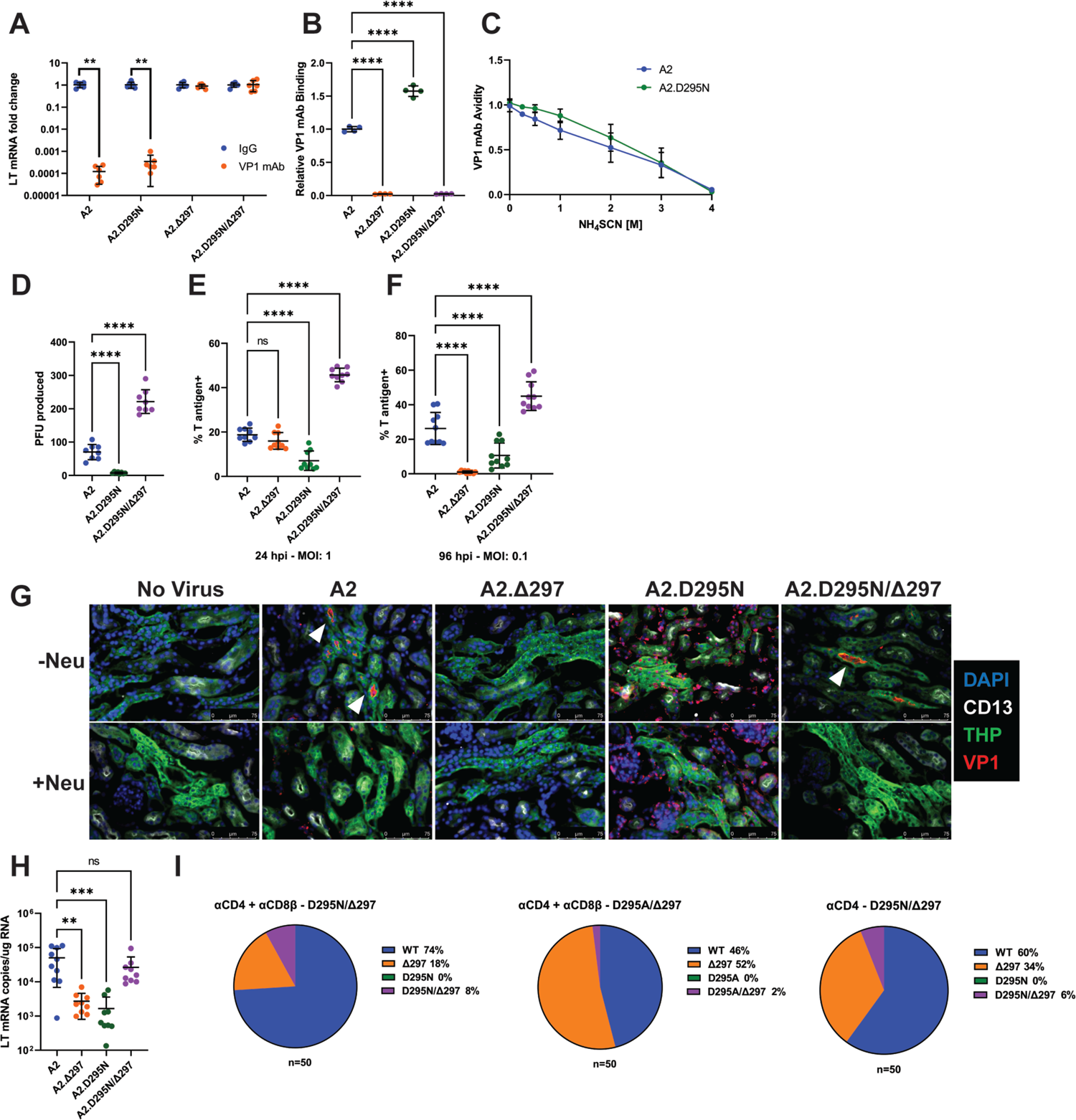
A compensatory mutation in VP1 arises to rescue defects in receptor binding caused by an Ab escape mutation. A. VP1 mAb neutralization assay with D295N and Δ297 VP1 mutant viruses. Viruses were preincubated with 10μg VP1 mAb or control IgG for 30 min prior to addition to 1 x 10^5^ NMuMG epithelial cells. A2 was diluted to an MOI of 0.1 PFU/cell, mutant viruses were diluted to match A2 by g.e. Viral LT mRNA levels were quantified 24 hpi and normalized for each virus to infection with control IgG (n=6). B. Binding of VP1 mAb to WT or VP1 mutant viruses. Wells were coated with 1 x 10^9^ g.e. of WT or VP1 mutant virus and incubated with VP1 mAb. VP1 mAb binding was quantified using an anti-rat secondary antibody and values were normalized to binding to WT virus (n=4). C. Binding avidity of VP1 mAb for A2 and A2.D295N. VP1 mAb binding to A2 and A2.D295N was performed as in B. Prior to detection of mAb binding, virus/mAb complexes were treated with varying concentrations of NH_4_SCN for 15 minutes. Binding at each concentration was normalized to binding at 0 M NH_4_SCN for each virus (n=4). D. Quantification of virus production in a single round of replication by plaque assay. Virus was added to 1 x 10^5^ A31 cells at an MOI of 0.1 PFU/cell. Cells were lysed at 60 hpi and infectious virus was quantified by plaque assay and divided by the input virus quantity (n=8). E. Frequency of T antigen positive cells 24 hpi with WT or mutant viruses. 1 x 10^5^ A31 cells were infected with A2 at an MOI of 1 PFU/cell or mutant viruses matched by g.e. Cells were collected at 24 hpi, permeabilized, stained for T ag protein, and quantified by flow cytometry. F. Frequency of T antigen positive cells 96 hpi with WT or mutant viruses. 1 x 10^5^ A31 cells were infected with A2 at an MOI of 0.1 PFU/cell or mutant viruses matched by g.e. Cells were collected at 96 hpi, permeabilized, stained for T ag protein, and quantified by flow cytometry. G. Detection of virus binding in kidney sections. PFA-fixed frozen kidney sections were treated with neuraminidase or buffer alone prior to incubation with WT or VP1 mutant virus. Sections were then stained for VP1 (red) and kidney markers [CD13 (white), THP (green)]. Neu: Neuraminidase. Representative of three independent experiments. H. LT mRNA levels in the kidney 4 dpi with WT or mutant viruses. WT mice were infected i.v. with 1 x 10^6^ PFU of A2 or mutant viruses matched by g.e. Viral LT mRNA levels were quantified by qPCR and compared to a standard curve (n=9-10). I. Detection of D295 and H297 mutations in the kidney. VP1 sequences were PCR amplified from kidney DNA samples of the mice that developed the D295N/Δ297 and D295A/Δ297 double mutant viruses in Figure 3A. VP1 clones were sequenced and screened for the presence of mutations at D295 and H297; the frequency of each mutation in 50 clones is shown. Error bars are mean ± SD. Data are from at least two independent experiments. Data were analyzed by Mann-Whitney U test with Holm-Šídák correction for multiple comparisons (A), one-way ANOVA with Dunnett’s multiple comparisons test (B, D-F), and Kruskal-Wallis test with Dunn’s multiple comparisons test (H). **p < 0.01, ***p < 0.001, ****p < 0.0001. See also **Figure S2**.

We next asked whether these mutations affected receptor usage. Parental A2 and A2.D295N/Δ297 showed preferential hemagglutination (HA) binding at acidic pH (**Figure S2C**). A2.D295N and A2.Δ297 exhibited poor HA activity, with A2.Δ297 showing no HA activity across the pH spectrum. As HA activity is dependent on sialic acid binding, we next examined the effect of neuraminidase pretreatment on virus binding to host cells. Binding by A2, A2.Δ297, and A2.D295N/Δ297 was dependent on sialyated host cell receptors, but binding by A2.D295N was refractory to neuraminidase treatment (**Figure S2D**). Despite these differences in binding, neuraminidase pretreatment caused a significant reduction 24 hpi in LT mRNA levels with low MOI infection and T ag^+^ cells with high MOI infection, indicating that infection by all viruses relied predominantly on a sialic acid-dependent pathway (**Figure S2E-F**). Furthermore, A2.Δ297 single mutant showed a drastic sensitivity to neuraminidase pretreatment, with 10,000-fold reduction in mRNA production, consistent with poor receptor binding and the lack of HA activity.

We next assessed how these mutations affected receptor binding and infection in the kidney. We incubated naïve kidney sections with virus and then stained for VP1 and kidney markers for proximal (CD13^+^) and distal (THP^+^) tubules (Baer et al., 1997; Tokonami et al., 2018). Parental A2 bound to the lumen of the distal convoluted tubules (DCT) and binding was abrogated by neuraminidase pretreatment (**Figure 5G**). Similar to A2, the A2.D295N/Δ297 showed specific, neuraminidase-sensitive binding to the DCT. A2.Δ297, however, showed no detectable binding in the kidney, whereas A2.D295N showed substantial binding to the abluminal side of the tubules but little-to no-binding to the luminal side. A2.D295N binding in the kidney was insensitive to neuraminidase treatment. Based on LT mRNA levels in the kidney at 4 dpi, both single mutant viruses were attenuated compared to A2, whereas A2.D295N/Δ297 showed similar levels of infection to A2 (**Figure 5H**). Together, these findings indicate that the Δ297 mutation conferred resistance to 8A7H5 neutralization, but impaired viral spread and infection within the kidney. The D295N mutation alone caused aberrant receptor binding and impaired infection *in vivo*, but when combined with Δ297 restored proper receptor binding in the kidney during early primary infection.

Based on the effects of the D295N and Δ297 mutations, we hypothesized that Ab escape by the Δ297 mutation was the initial driver of mutant virus emergence *in vivo*, with the emergence of the mutation at D295 occurring secondary to compensate for the defect in virulence. To test this, we cloned and sequenced VP1 from the kidney of the T cell-depleted mice that developed the D295N/Δ297 and D295A/Δ297 double mutant viruses. We identified sequences containing WT VP1, Δ297, and D295N/Δ297 or D295A/Δ297, but not sequences containing the D295N/A mutations alone (**Figure 5I**). The absence of single D295 mutant sequences indicates the order of mutation emergence was Δ297 followed by D295N/A.

### Heightened neurovirulence by a VP1 double mutation virus

Given the increased replication of A2.D295N/Δ297 *in vitro*, we next investigated whether this virus showed altered kidney pathology in an immunocompromised host. We infected μMT mice intravenously (i.v.) with A2 or A2.D295N/Δ297, which resulted in chronic viremia 30 dpi by each virus (**Figure 6A**). Kidneys from A2-infected mice had foci of lymphocytic inflammation within the cortical interstitium that extended into the tubular epithelium (**Figure 6B**). In marked contrast, kidneys A2.D295N/Δ297-infected mice had only small, scattered collections of mononuclear cells. Consistent with these histopathologic differences, A2-infected kidneys also exhibited numerous, large VP1^+^ foci, but those in A2.D295N/Δ297-infected kidneys were far fewer and smaller (**Figure 6C-D**). To determine if A2.D295N/Δ297 was neurovirulent, we infected WT mice intracranially (i.c.) with either A2 or A2.D295N/Δ297 virus. Four dpi, A2-infected brains showed only sparse VP1^+^ ependymal cells lining the ventricles. In contrast, brains of A2.D295N/Δ297-infected mice had extensive VP1^+^ ependymal cells (**Figure 6E**).

**Figure 6.**
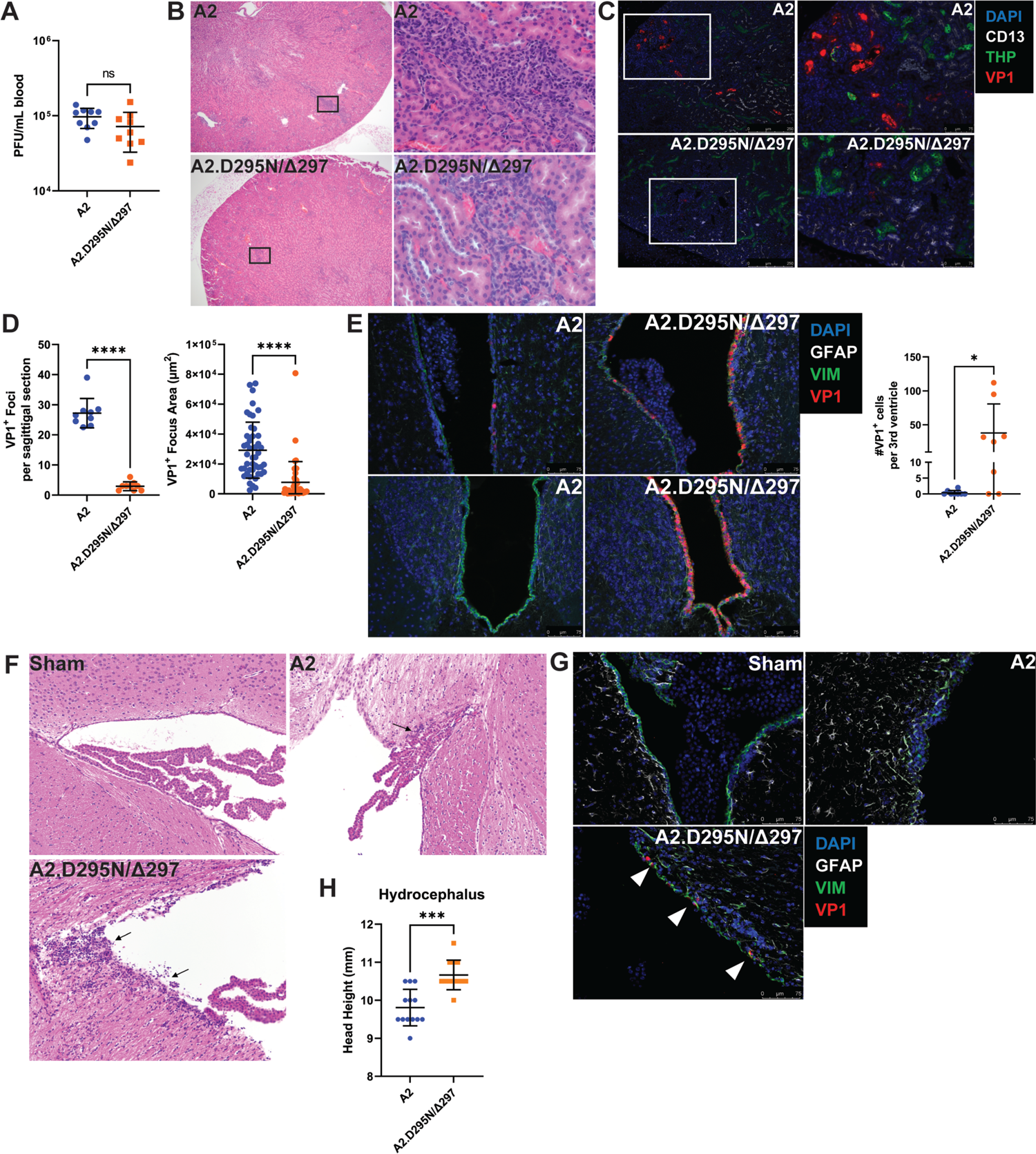
Lower kidney tropism and heightened neurovirulence by a VP1 double mutant virus. A. Viremia in μMT mice 30 dpi with A2 or A2.D295N/Δ297 i.v. Viremia was quantified by plaque assay from whole blood (n=9). B. H&E stained sagittal sections of kidneys from μMT mice 30 dpi after i.v. inoculation with A2 or A2.D295N/Δ297. Left: 50x magnification. Right: 500x magnification. C. Foci of VP1^+^ cells in kidneys from μMT mice 30 dpi with A2 or A2.D295N/Δ297 i.v. Kidneys were stained for CD13 (white), THP (green) and VP1 (red). D. Quantification of the number (n=9) and size (n=42-45) of VP1^+^ foci in kidneys from μMT mice 30 dpi with A2 or A2.D295N/Δ297 i.v. Foci number is the average of two sagittal kidney sections per mouse. For quantifying foci area, five random foci per mouse or the maximum number of foci found were imaged and the area of each VP1^+^ focus was calculated using ImageJ. E. VP1^+^ cells in the lateral (top) and third (bottom) ventricles of WT mice 4 dpi with A2 or A2.D295N/Δ297 inoculated i.c. and quantification of VP1^+^ cells in the 3^rd^ ventricle (n=8). F. H&E stained coronal sections from brains of sham (top left), A2 (top right), or A2.D295N/Δ297 (bottom left) i.c.-inoculated mice 30 dpi (200x magnification). Arrows indicate sites of ependymal inflammation. G. VP1 staining in the ventricles of WT mice 30 dpi with A2 or A2.D295N/Δ297 i.c. VP1^+^ cells are indicated with white markers. H. Quantification of hydrocephalus 30 dpi with A2 or A2.D295N/Δ297 after i.c. inoculation. Coronal head height was measured with a Vernier caliper in line with the ear canal to the nearest 0.5 mm (n=12-13). Error bars are mean ± SD. Data are from at least two independent experiments. Data were analyzed by Mann-Whitney U test (A, D, E, and H). *p < 0.05, ***p < 0.001, ****p < 0.0001.

Consistent with this dramatic difference in extent of ependymal infection, A2-infected brains at 30 dpi had only small ependymal lymphocytic aggregates, whereas A2.D295N/Δ297-infected brains contained more extensive ependymitis as well as periventricular edema (**Figure 6F**). Additionally, of the sections examined only brains from mice persistently infected with A2.D295N/Δ297 had VP1^+^ cells in the periventricular region (**Figure 6G**). Hydrocephalus is consistently seen in mice i.c. inoculated with A2 MuPyV (Lauver et al., 2020; Mockus et al., 2020). A2.D295N/Δ297-infected mice developed hydrocephalus to a significantly higher degree than A2-infected mice (**Figure 6H**). These data show that the endogenously derived A2.D295N/Δ297 VP1 mutant virus had lower tropism for the kidney, but dramatically higher capacity to infect the cerebral ventricular system than the parental A2 virus. Similarly, impaired kidney pathogenesis but retained neurovirulence is a hallmark feature of A2 virus carrying the VP1 V296F mutation, sequence-equivalent to the frequent S268F VP1 mutation in JCPyV-PML (Lauver et al., 2020). Thus, VP1 neutralizing Ab in T cell-deficient hosts can select rare Ab-escape virus variants possessing high neuropathogenicity.

## Discussion

Although depressed anti-JCPyV T cell immunity is the dominant risk factor for PML, JCPyVs in the CNS of PML patients have VP1 mutations that allow evasion of antiviral (Cortese et al., 2021; Jelcic et al., 2015; Lauver et al., 2020; Pavlovic et al., 2018; Ray et al., 2015). To connect T cell deficiency and VP1-specific Ab-escape JCPyV variants, we investigated T cell insufficiency in mice infected by MuPyV in the setting of a restricted VP1 antibody response.

Our findings allowed us to develop the following model integrating T cell and humoral deficiencies with emergence of neurovirulent PyV. First, impaired T cell control results in PyV resurgence in the kidney, the central reservoir for persistent PyV infection. Elevated viral replication is accompanied by low-level, stochastic mutagenesis of the viral dsDNA genome, where particular mutations in VP1 abrogate binding by VP1-specific Ab. By extension, the host must have an inherited or acquired VP1 Ab response that targets few VP1 epitopes. A subset of viruses carrying Ab-escape VP1 mutations acquire the capacity to replicate in the CNS. T cell deficiency further handicaps a second line of defense by antiviral T cells to control Ab-escape VP1 variant viruses. In summary, T cell deficiency acts at two levels to set the stage for emergence of antibody-escape VP1 mutations endowed with neurotropic potential. The kidney is the dominant site of JCPyV persistence (Berger et al., 2017). We observed preferential binding of MuPyV to the DCT, mirroring what has been reported for JCPyV, and virus replication in the DCT following T cell loss (**Figures 1D and 5G**)(Haley et al., 2015).

Virus binding and replication within the DCT may provide a level of local protection from neutralizing antibodies, necessitating T cell control to limit virus replication within the local kidney environment. Conditions of immune suppression deprive the kidney of this cellular immune control over PyV infection, leading to virus resurgence there. Consistent with this are reports of depressed JCPyV-specific T cell responses in individuals treated with natalizumab and increased JCPyV shedding in the urine of immunosuppressed individuals (Behzad-Behbahani et al., 2004; Chen et al., 2009; Delbue et al., 2015; Perkins et al., 2012). T cells are the predominant cell type responsible for the production of interferon gamma (IFN-γ), which limits BKPyV replication in kidney cells and is necessary for effective control of MuPyV kidney infection (Abend et al., 2007; Byers et al., 2007; Wilson et al., 2011). T cell loss during PyV infection thus eliminates the major source of IFN-γ and virus control.

PML is associated with conditions of CD4 T cell impairment, including HIV/AIDS and idiopathic CD4 T cell lymphopenia (Berger et al., 1987, 1998; Pavlovic et al., 2018). Decreases in CD4, but not CD8, T cells are seen in the CSF of natalizumab-treated MS patients and CD4 T cell epitope-escape JCPyVs have been found in PML patients (Jelcic et al., 2016; Schneider-Hohendorf et al., 2014; Stüve et al., 2006). We observed a higher kidney virus levels at 10 days post depletion and a higher frequency of viremia in CD4 T cell-depleted mice than CD8T cell-depleted mice (**Figures 1C and 2B**). CD4 T cells can exert direct antiviral effector activities as well as provide help to maintain optimal virus-specific CD8 T cell responses in nonlymphoid organs, including the brain and lung (Ren et al., 2020; Son et al., 2021). Tissue-resident CD4 T cells have also been shown to promote development of protective B cell responses in the lungs of influenza virus-infected mice (Son et al., 2021). It will be important to elucidate how CD4 T cells prevent PyV resurgence in the kidneys and viremia by Ab-escape variant viruses.

Utilizing the host DNA replication machinery, PyVs are regarded as having highly stable genomes with a low incidence of mutations (Sanjuán and Domingo-Calap, 2016). Clinical and experimental data, however, indicate that mutations arise during the course of PyV infection. JCPyV and BKPyV isolates from PML and BKPVN patients, respectively, are characterized by mutations in both the VP1 capsid protein and the non-coding control region (NCCR) (Boldorini et al., 2009; Gorelik et al., 2011; Krautkrämer et al., 2009; Peretti et al., 2018; Reid et al., 2011; Tremolada et al., 2010; Vaz et al., 2000; Zheng et al., 2005b, 2005a). Likewise, rearrangements in the NCCR of BKPyV emerge after extended passage *in vitro*, and VP1 mutations arise in BKPyV and MuPyV after serial passage in the presence of neutralizing mAb (Lauver et al., 2020; Lindner et al., 2019; Zhao and Imperiale, 2021). These data suggest that conditions favoring certain mutations (e.g., mutations enhancing replication or enabling antibody evasion) can promote the emergence and outgrowth of viruses with particular genomic alterations.

Several mechanisms provide avenues for VP1 mutations. Missense mutations may represent failed attempts by the host cell to restrict viral replication. The APOBEC3 family of proteins are cytosine deaminases that induce mutagenic damage in the genomes of DNA viruses as a mechanism of antiviral defense. APOBEC3B is upregulated during BKPyV nephritis and has been implicated in the emergence of BKPyV VP1 mutations in kidney transplant recipients and mutations in the Merkel cell PyV genome (Peretti et al., 2018; Que et al., 2021). The presence of deletion mutations could represent the remnants of double stranded breaks repaired by nonhomologous end joining (NHEJ). DNA double strand breaks occur during PyV infection, and aspects of the DNA damage response pathway are required for efficient PyV replication (Erickson and Garcea, 2019; Heiser et al., 2016; Jiang et al., 2012; Justice et al., 2019; Sowd et al., 2013). The NCCR of BKPyV undergoes significant recombination events resulting from NHEJ during replication *in vitro* (Zhao and Imperiale, 2021). It is possible that NHEJ repair of breaks in VP1 during replication results in a variety of recombinations and small deletions. Those that are tolerated by the virus and confer a selective advantage, in this case in-frame deletions conferring Ab escape, are thus able to emerge and be detected.

JCPyV VP1 mutations identified in PML patients are associated with impaired receptor binding and infectivity and only archetype JCPyV is found in the urine of PML patients, despite the presence of VP1 mutant virus in the blood and CSF of these patients (Geoghegan et al., 2017; Gorelik et al., 2011; Maginnis et al., 2013; Reid et al., 2011; Zheng et al., 2005a). Here, we find that the selective pressure of antibody escape drove outgrowth of VP1 mutants with varying kidney infectivity, implicating immune evasion, rather than tropism, as the primary driver of VP1 mutations under these conditions (**Figure 3C-D**). The initial emergence of the Δ297 mutation, despite severe defects in kidney tropism, indicates that Ab escape can promote the outgrowth of otherwise severely impaired PyV variants, a feature seen in several other chronic viral infections (Kalinina et al., 2003; Kinchen et al., 2018; Lynch et al., 2015).

The defect in spread and infectivity *in vivo* by the Δ297 mutation readily selected for a variety of secondary point mutations that restored or even enhanced virulence, generating viruses able to efficiently spread *in vitro*. The point mutations found with the Δ297 mutation all involved a positive net change in charge (**Figure 3A**). These mutations are all located near the receptor binding pocket, and most of the mutated side chains face toward the location of receptor binding. However, with the exception of E91, these residues have not been reported to be involved in receptor binding (Buch et al., 2015; Stehle and Harrison, 1997). Mutation of E91 to glycine causes impaired infection by promoting binding to nonproductive pseudoreceptors (Bauer et al., 1999; Buch et al., 2015; Qian and Tsai, 2010). Similarly, the D295N mutation alone caused aberrant pseudoreceptor binding that impaired infectivity *in vitro* and *in vivo* (**Figures 5E and 5H**). In the context of the Δ297 mutation however, these normally deleterious mutations synergized to restore proper receptor binding and even enhance virulence. The D295N/Δ297 double mutant displayed increased infectivity and virus production *in vitro* (**Figure 5D-F**). *In vivo,* the double mutant showed decreased kidney virulence but increased acute infection of the ependyma and evidence of heightened chronic infection in the periventricular region, as well as increased CNS pathology (**Figure 6E-G**). Retention of neurotropism and a loss of kidney tropism was previously seen with the V296F MuPyV VP1 mutant and may a common feature of PML-associated mutations (Lauver et al., 2020). This increased CNS virulence could stem from more efficient receptor binding and release, leading to increased viral dissemination throughout the ventricular system and resulting in elevated infection of the ependyma during persistence.

Increased spread and pathogenesis resulting from altered receptor binding is seen with a separate MuPyV HI loop mutation, V296A (Bauer et al., 1995, 1999; Buch et al., 2015). As both the D295 and H297 residues face into the pore at the center of VP1, the mutations may facilitate more efficient interactions with the VP2/3 minor capsid proteins, the C-termini of which extend into the pore (Chen et al., 1998).

PML patients typically carry a mutant JCPyV with a single VP1 mutation, rather than the double mutants we saw emerge in several mice (**Table S1**)(Gorelik et al., 2011; Reid et al., 2011). This difference may stem from the nature of Ab escape mutation, with substitution to a bulkier side chain in JCPyV (e.g. L55F, S266F/Y, and S268F/Y) rather than the deletion of H297 seen in MuPyV. Although both types of mutations alter/impair infectivity, the bulky substitutions in JCPyV may do so by occluding a portion of the receptor binding pocket, which there is a reduced opportunity for a secondary mutation to compensate for. Already the smallest external loop in VP1, the HI loop of JCPyV (M261 to S268: 8 residues) is one residue shorter than the HI loop of MuPyV (W288 to V296: 9 residues)(Liddington et al., 1991). Thus, the HI loop of JCPyV may be too short to tolerate even a single amino acid deletion and still allow proper position of the H and I β strands.

Insufficient humoral control of PyV mutants opens numerous pathways for viral invasion of the CNS. The lack of neutralization could promote infection of susceptible immune cells in the blood and subsequent trafficking to the brain (Dubois et al., 1997; Wollebo et al., 2015). Alternatively, the absence of neutralizing antibody could enable virus within the vasculature of the brain to infect the cells of the blood-brain barrier (BBB) or blood-CSF barrier (BCSFB). Brain microvascular endothelial cells are susceptible to JCPyV infection, and could provide viral access into the CNS parenchyma (Chapagain et al., 2007). The BCSFB, formed by the choroid plexus in the brain ventricles, is another possible route of entry by the virus into the brain. JCPyV binds choroid plexus epithelium and infects choroid plexus epithelial cells (Corbridge et al., 2019; Haley et al., 2015; O’Hara et al., 2018, 2020). Infection of choroid plexus epithelium would enable virus spread into the CSF, dissemination throughout the ventricular system, and access to the brain parenchyma via infection of the ependymal cells lining the ventricles. In line with this possibility is the permissivity of ependymal cells for productive MuPyV infection (Lauver et al., 2020; Mockus et al., 2020).

Our findings demonstrate a “2-hit” requirement for development of VP1 mutant PyVs: (1) a narrow VP1 antibody response; and (2) T cell insufficiency, which acts both to limit virus replication in the kidney and eliminate VP1 variants that evade antiviral antibodies. T cell loss leads to virus resurgence that in the context of a narrow VP1 antibody response results in viremia by Ab-escape mutant viruses. Among these Ab-escape variants will be those carrying the potential for neurovirulence. The stochastic evolution of VP1 mutations, together with depressed anti-PyV T and B cell immunity, may account for the long timeframe between PML and iatrogenic immunosuppression, the array of PML-associated agents with unrelated mechanisms of action, and the rarity of PML in susceptible hosts.

## Acknowledgements

We thank the staff of the Penn State College of Medicine Flow Cytometry Core Facility for assistance with flow cytometry experiments; the Comparative Medicine Histology Core for sample processing; Kimberly Erickson and Robert Garcea for the generous gifts of VP1 pentamers and rabbit VP1 antisera; and the staff of the Penn State College of Medicine Department of Comparative Medicine. This work was supported by NIH grants 5R01NS088367 and 5R01NS092662.

## Author contributions

Conceptualization, M.D.L. and A.E.L.; methodology, M.D.L. and A.E.L.; investigation, M.D.L., G.J., K.N.A., C.S.S., C.S.A., and S.N.C; writing – original draft, M.D.L. and A.E.L.; writing – review & editing, M.D.L. and A.E.L.; funding acquisition, A.E.L.

---

**Supplemental Figure 1. VP1 mAb injections give μMT mice Ab-escape blind spots, Related to Figure 1.** A. VP1-specifc IgG 30 dpi from WT or μMT mice detected by ELISA with VP1 pentamers (n=10-12). B. Viremia at 30 dpi in WT, μMT, or μMT mice treated with VP1 mAb starting 4 dpi. LOD: Limit of detection (n=8). C-D. Neutralization of A2 (C) or A2.Δ295 (D) by sera from WT or VP1-mAb treated μMT mice 20 dpi. LT mRNA fold change is relative to infection by each virus in the absence of serum (n=8-9). E. Neutralizing titers against A2 and A2.Δ295 from WT or VP1-mAb treated μMT mice 20 dpi. ND: Not determined (n=8-9). Error bars are mean ± SD. Data are from at least two independent experiments.

**Supplemental Figure 2. Defects in mutant virus spread and receptor binding, Related to Figure 5.** A. Plaque formation by WT and mutant viruses in A31 fibroblasts 6dpi. B. Encapsidated genomes produced by WT and mutant viral DNA transfection. Data is relative to WT (n=9). C. pH-dependent hemagglutination curves for WT and mutant viruses. The reported HA titer was the highest dilution showing hemagglutination (n=3). D. Percent of virus binding lost with neuraminidase pre-treatment. Bound virus was detected with polyclonal VP1 Ab and quantified by flow cytometry. For each virus, the percent of binding lost to neuraminidase treatment was calculated by comparing virus binding with and without neuraminidase treatment (n=6). E. Quantification of T antigen positive cells 24 hpi with WT or mutant viruses with neuraminidase pretreatment. Cells were treated and infected as in (E). At 24 hpi, cells were collected, permeabilized, stained for T ag protein, and quantified by flow cytometry (n=6). F. Quantification of LT mRNA levels with neuraminidase pretreatment. A31 cells were treated with neuraminidase and then infected with A2 at an MOI of 0.1 PFU/cell or mutant viruses matched by g.e. Viral LT mRNA levels were quantified by qPCR and compared to a standard curve (n=6). Error bars are mean ± SD. Data are from at least two independent experiments. Data were analyzed by one-way ANOVA with Dunnett’s multiple comparisons test (B and D) or Mann-Whitney U test with Holm-Šídák correction for multiple comparisons (E-F). **p < 0.01, ***p < 0.001.

## Materials and Methods

### Mice

C57BL/6 mice were purchased from the National Cancer Institute and μMT mice were purchased from Jackson Laboratories. Mice were housed and bred under specific pathogen-free conditions. Male and female mice 6-12 weeks of age were used for experiments. All mouse experiments were approved by the Penn State College of Medicine Institutional Animal Care and Use committee.

### Cell lines

NMuMG and BALB/3T3 clone A31 cell lines were purchased from ATCC. Cell lines were authenticated by STR profiling (ATCC), mycoplasma negative, used at low passage, and examined for correct morphology. The 8A7H5, H35-17.2, and GK1.5 hybridomas were grown in PFHM-II Protein-Free Hybridoma Medium (Thermofisher) (Dialynas et al., 1983; Golstein et al., 1982; Swimm et al., 2010). mAb was produced in CELLine bioreactor flasks (Corning). All other cells were kept in Dulbecco’s Minimal Eagle Media supplemented with 10% fetal bovine serum, 100 U/mL penicillin, and 100 U/mL streptomycin.

### Viruses

All experiments were done using the A2 strain of MuPyV. Viral stocks were generated by transfection of viral DNA into NMuMG cells. Mutant viruses were generated by site-directed mutagenesis with forward and reverse primers for each mutation (**Table S2**). DNA was isolated from the virus stocks and sequenced to confirm the presence of the mutation. The A2.Δ294, A2.Δ295, and A2.V296F viruses were generated previously (Lauver et al., 2020).

### Virus titering and sequencing

Viruses were titered by plaque assay on A31 fibroblasts or by qPCR for encapsidated genomes (Lukacher and Wilson, 1998). For genome titering, 1 μL of virus lysate was treated with 250 U of benzonase nuclease (Sigma) at 37°C for 1 h. Viral genomes were isolated using the Purelink Viral RNA/DNA mini kit and genome ratios were determined by Taqman qPCR (Wilson et al., 2012).

### Mouse infections and treatments

Mice were infected via the hind footpad with 1 x 10^6^ PFU of A2 MuPyV. Challenge infections with A2.Δ295 were given with 1 x 10^3^ PFU i.v. For comparisons of mutant viruses, mice were infected i.v. with 1 x 10^6^ PFU of A2 or the mutant virus matched by g.e. For comparisons of brain infection, mice were infected i.c. with 5×10^5^ PFU of A2 or A2.D295N/Δ297 matched by g.e. µMT mice were injected i.p. weekly with 250 µg of 8A7H5 starting 4 dpi. For T cell depletions, mice were injected i.p. weekly with 250 µg of GK1.5 and H35-17.2 or control IgG.

### Virus infections in vitro

A31 or NMuMG cells were seeded in 12-well plates at density of 5×10^4^ cells/well the day before infection. Cells were washed with Iscove’s media with 0.1% BSA prior to infection with the specified MOI of A2 or mutant virus matched by g.e. Infections were performed at 4°C for 1.5 hours; unbound virus was then washed out and the cells were returned to DMEM with 10% FBS. For infections with neuraminidase pretreatment, cells were incubated with or without *Vibrio cholerae* neuraminidase (Sigma 1:200) in neuraminidase buffer (PBS with 1mM CaCl_2_, 1mM MgCl_2_) at 37°C for 30 min. Neutralization assay infections were performed at an MOI of 0.1 PFU/cell with A2 virus or VP1 mutant virus matched by g.e. For serum neutralization assays, virus was incubated with the indicated serial dilution of serum at 4°C for 30 min prior to addition to cells. For neutralization assays with VP1 mAb, virus was incubated with 10 µg of 8A7H5 or control IgG at 4°C for 30 min prior to addition to cells. For quantification of encapsidated genome production, A31 cells were transfected with equal amounts of WT or mutant viral DNA using Lipofectamine 3000 (ThermoFisher). At 24 hours the media was removed, and the cells were washed and placed in fresh media to remove free DNA. At 72 hours, the cells and media were collected and the amount of encapsidated genomes were quantified as above. For images of plaque formation, plaque assays were imaged 6dpi with an Olympus IX73 inverted microscope with a QImaging Retiga 6000 Mono camera.

### Viral mRNA and DNA quantification

Total RNA was isolated with TRIzol Reagent (Thermofisher) and phenol:chloroform extraction. Total cDNA was generated from 1-2 µg of RNA using random hexamer primers and Revertaid RT (Thermofisher). Taqman qPCR was used to quantify LT mRNA levels with normalization to TATA-Box Binding protein and compared to a standard curve to determine copy number (Maru et al., 2017). DNA was isolated with the Wizard Genomic DNA Purification Kit (Promega). Viral DNA was quantified by Taqman qPCR and compared to a standard curve to determine copy number (Wilson et al., 2012).

### TA cloning for VP1 sequencing

VP1 sequences were PCR amplified from kidney DNA and cloned using the TOPO TA cloning kit (Thermofisher). Clones were screened by restriction digest for the presence of a VP1 sequence and sequenced. VP1 sequences were screened from the presence of the D295A/N and Δ297 mutations.

### Hemagglutination Assay

Viruses were diluted to 1 x 10^7^ g.e./μL then serially two-fold diluted in PBS at the indicated pH. Virus dilutions were combined 1:1 with 0.45% sheep erythrocytes (Innovative Research) and incubated overnight at 4°C. The highest dilution at each pH showing hemagglutination was reported as the HA titer.

### ELISA

Full-length MuPyV VP1 pentamers were kindly provided by Robert Garcea (University of Colorado, Boulder). ELISA wells were coated overnight at 4°C with 50 ng of VP1 pentamer or 1 x 10^7^ PFU of A2 or g.e. matched mutant virus. For avidity measurements, 8A7H5-virus complexes were treated with NH_4_SCN for 15 min before the addition of the secondary and detection. 8A7H5 binding was normalized for each virus to signal in the absence of NH_4_SCN (Lauver et al., 2020; Pullen et al., 1986).

### Flow cytometry

T cell depletions were confirmed in the peripheral blood at euthanasia by staining with antibodies for CD45 (30-F11), CD8α (53-6.7), and CD4 (RM4-5) (Biolegend, 1:200). For quantification of *in vitro* infections, cells were trypsinized and stained with Fixable Viability Dye (ThermoFisher, 1:1000) followed by treatment with eBiosience Fixation/Permeabilization reagent (ThermoFisher). Cells were then stained with rat polyclonal T antigen antibody (1:500) followed by an anti-rat secondary (Biolegend, 1:200). For measuring sialic acid binding dependence, trypsinized cells were treated for 30 min at 37°C in the presence or absence of *Vibrio cholerae* neuraminidase (Sigma, 1:200). Bound virus was detected with rabbit polyclonal VP1 antibody (1:10000) followed by an anti-rabbit secondary (Biolegend, 1:200). Samples were acquired on an LSRFortessa flow cytometer (BD Biosciences) and analyzed using FlowJo software (Tree Star).

### Immunofluorescence and histological imaging

Kidneys were immersion-fixed in neutral buffered formalin (NBF) overnight prior to processing and paraffin-embedding. For brain preparation, mice were perfused with NBF and whole heads were fixed overnight in NBF. The brains were then removed for processing and embedding.

Formalin-fixed paraffin embedded kidney and brain sections were stained with VP1 (1:1000), CD13 (Abcam, 1:500), GFAP (Abcam, 1:1000), Vimentin (R&D, 1:500) and THP (R&D, 1:1000) antibodies. Hematoxylin and eosin (H&E) stained sagittal sections of kidneys and coronal sections of brains were evaluated by a renal pathologist and a neuropathologist, respectively, in blinded fashion. For virus binding to kidney sections, paraformaldehyde-fixed kidneys were embedded in Tissue-Tek O.C.T. Compound (Sakura) and cyrosectioned. Sections were treated with *Vibrio cholerae* neuraminidase (Sigma, 1:100) or buffer for 30 min at 37°C and then incubated with virus lysate for 1.5 h. Sections were then stained with VP1 (1:10000), CD13 (Abcam, 1:500), and THP (R&D, 1:1000) antibodies. Secondary antibodies used were anti-rabbit Alexa Fluor 647 (Jackson Immunoresearch, 1:500), anti-goat Alexa Fluor 488 (Jackson Immunoresearch, 1:500) and Alexa Fluor 555 anti-rat secondary (Abcam, 1:500).

Samples were mounted with Prolong Gold Antifade Mountant with DAPI (ThermoFisher). Samples were imaged on a Leica DM4000 fluorescence microscope. For representative fluorescence images, adjustments for brightness/contrast were done uniformly to all images in the group using LAS X (Leica).

### Statistical Analysis

All data are displayed as mean ± SD. The statistical tests performed are listed with the respective figures and were performed using Prism software (Graphpad). p values of ≤0.05 were considered significant. Statistical methods were not used to pre-determine sample sizes. Sample sizes represent individual mice or biological replicates. VP1^+^ foci were counted in a blinded fashion, blinding was not employed for other experiments. All sample sizes and number of repeats are included in the Figure Legends.

## References

1. Abend, J.R., Low, J.A., and Imperiale, M.J. (2007). Inhibitory effect of gamma interferon on BK virus gene expression and replication. J. Virol. 81, 272–279.

2. Baer, P.C., Nockher, W.A., Haase, W., and Scherberich, J.E. (1997). Isolation of proximal and distal tubule cells from human kidney by immunomagnetic separation. Technical note. Kidney Int. 52, 1321– 1331.

3. Bar, K.J., Sneller, M.C., Harrison, L.J., Justement, J.S., Overton, E.T., Petrone, M.E., Salantes, D.B., Seamon, C.A., Scheinfeld, B., Kwan, R.W., et al. (2016). Effect of HIV Antibody VRC01 on Viral Rebound after Treatment Interruption. N. Engl. J. Med. 375, 2037–2050.

4. Bauer, P.H., Bronson, R.T., Fung, S.C., Freund, R., Stehle, T., Harrison, S.C., and Benjamin, T.L. (1995). Genetic and structural analysis of a virulence determinant in polyomavirus VP1. J. Virol. 69, 7925–7931.

5. Bauer, P.H., Cui, C., Liu, W.R., Stehle, T., Harrison, S.C., DeCaprio, J.A., and Benjamin, T.L. (1999). Discrimination between sialic acid-containing receptors and pseudoreceptors regulates polyomavirus spread in the mouse. J. Virol. 73, 5826–5832.

6. Behzad-Behbahani, A., Klapper, P.E., Vallely, P.J., Cleator, G.M., and Khoo, S.H. (2004). Detection of BK virus and JC virus DNA in urine samples from immunocompromised (HIV-infected) and immunocompetent (HIV-non-infected) patients using polymerase chain reaction and microplate hybridisation. J. Clin. Virol. Off. Publ. Pan Am. Soc. Clin. Virol. 29, 224–229.

7. Berger, J.R., and Fox, R.J. (2016). Reassessing the risk of natalizumab-associated PML. J. Neurovirol. 22, 533–535.

8. Berger, J.R., Kaszovitz, B., Post, M.J., and Dickinson, G. (1987). Progressive multifocal leukoencephalopathy associated with human immunodeficiency virus infection. A review of the literature with a report of sixteen cases. Ann. Intern. Med. 107, 78–87.

9. Berger, J.R., Pall, L., Lanska, D., and Whiteman, M. (1998). Progressive multifocal leukoencephalopathy in patients with HIV infection. J. Neurovirol. 4, 59–68.

10. Berger, J.R., Miller, C.S., Danaher, R.J., Doyle, K., Simon, K.J., Norton, E., Gorelik, L., Cahir-McFarland, E., Singhal, D., Hack, N., et al. (2017). Distribution and Quantity of Sites of John Cunningham Virus Persistence in Immunologically Healthy Patients: Correlation With John Cunningham Virus Antibody and Urine John Cunningham Virus DNA. JAMA Neurol. 74, 437–444.

11. Bloomgren, G., Richman, S., Hotermans, C., Subramanyam, M., Goelz, S., Natarajan, A., Lee, S., Plavina, T., Scanlon, J.V., Sandrock, A., et al. (2012). Risk of natalizumab-associated progressive multifocal leukoencephalopathy. N. Engl. J. Med. 366, 1870–1880.

12. Boldorini, R., Allegrini, S., Miglio, U., Paganotti, A., Veggiani, C., Mischitelli, M., Monga, G., and Pietropaolo, V. (2009). Genomic mutations of viral protein 1 and BK virus nephropathy in kidney transplant recipients. J. Med. Virol. 81, 1385–1393.

13. Buch, M.H.C., Liaci, A.M., O’Hara, S.D., Garcea, R.L., Neu, U., and Stehle, T. (2015). Structural and Functional Analysis of Murine Polyomavirus Capsid Proteins Establish the Determinants of Ligand Recognition and Pathogenicity. PLoS Pathog. 11, e1005104.

14. Byers, A.M., Hadley, A., and Lukacher, A.E. (2007). Protection against polyoma virus-induced tumors is perforin-independent. Virology 358, 485–492.

15. Caskey, M., Klein, F., Lorenzi, J.C.C., Seaman, M.S., West, A.P., Buckley, N., Kremer, G., Nogueira, L., Braunschweig, M., Scheid, J.F., et al. (2015). Viraemia suppressed in HIV-1-infected humans by broadly neutralizing antibody 3BNC117. Nature 522, 487–491.

16. Caskey, M., Schoofs, T., Gruell, H., Settler, A., Karagounis, T., Kreider, E.F., Murrell, B., Pfeifer, N., Nogueira, L., Oliveira, T.Y., et al. (2017). Antibody 10-1074 suppresses viremia in HIV-1-infected individuals. Nat. Med. 23, 185–191.

17. Chapagain, M.L., Verma, S., Mercier, F., Yanagihara, R., and Nerurkar, V.R. (2007). Polyomavirus JC infects human brain microvascular endothelial cells independent of serotonin receptor 2A. Virology 364, 55–63.

18. Chen, X.S., Stehle, T., and Harrison, S.C. (1998). Interaction of polyomavirus internal protein VP2 with the major capsid protein VP1 and implications for participation of VP2 in viral entry. EMBO J. 17, 3233– 3240.

19. Chen, Y., Bord, E., Tompkins, T., Miller, J., Tan, C.S., Kinkel, R.P., Stein, M.C., Viscidi, R.P., Ngo, L.H., and Koralnik, I.J. (2009). Asymptomatic reactivation of JC virus in patients treated with natalizumab. N. Engl. J. Med. 361, 1067–1074.

20. Corbridge, S.M., Rice, R.C., Bean, L.A., Wüthrich, C., Dang, X., Nicholson, D.A., and Koralnik, I.J. (2019). JC virus infection of meningeal and choroid plexus cells in patients with progressive multifocal leukoencephalopathy. J. Neurovirol. 25, 520–524.

21. Cortese, I., Reich, D.S., and Nath, A. (2021). Progressive multifocal leukoencephalopathy and the spectrum of JC virus-related disease. Nat. Rev. Neurol. 17, 37–51.

22. Delbue, S., Elia, F., Carloni, C., Pecchenini, V., Franciotta, D., Gastaldi, M., Colombo, E., Signorini, L., Carluccio, S., Bellizzi, A., et al. (2015). JC virus urinary excretion and seroprevalence in natalizumab-treated multiple sclerosis patients. J. Neurovirol. 21, 645–652.

23. Dialynas, D.P., Quan, Z.S., Wall, K.A., Pierres, A., Quintáns, J., Loken, M.R., Pierres, M., and Fitch, F.W. (1983). Characterization of the murine T cell surface molecule, designated L3T4, identified by monoclonal antibody GK1.5: similarity of L3T4 to the human Leu-3/T4 molecule. J. Immunol. Baltim. Md 1950 131, 2445–2451.

24. Dubois, V., Dutronc, H., Lafon, M.E., Poinsot, V., Pellegrin, J.L., Ragnaud, J.M., Ferrer, A.M., and Fleury, H.J. (1997). Latency and reactivation of JC virus in peripheral blood of human immunodeficiency virus type 1-infected patients. J. Clin. Microbiol. 35, 2288–2292.

25. Erickson, K.D., and Garcea, R.L. (2019). Viral replication centers and the DNA damage response in JC virus-infected cells. Virology 528, 198–206.

26. Fox, R.J., and Rudick, R.A. (2012). Risk stratification and patient counseling for natalizumab in multiple sclerosis. Neurology 78, 436–437.

27. Geoghegan, E.M., Pastrana, D.V., Schowalter, R.M., Ray, U., Gao, W., Ho, M., Pauly, G.T., Sigano, D.M., Kaynor, C., Cahir-McFarland, E., et al. (2017). Infectious Entry and Neutralization of Pathogenic JC Polyomaviruses. Cell Rep. 21, 1169–1179.

28. Golstein, P., Goridis, C., Schmitt-Verhulst, A.M., Hayot, B., Pierres, A., van Agthoven, A., Kaufmann, Y., Eshhar, Z., and Pierres, M. (1982). Lymphoid cell surface interaction structures detected using cytolysis-inhibiting monoclonal antibodies. Immunol. Rev. 68, 5–42.

29. Gorelik, L., Reid, C., Testa, M., Brickelmaier, M., Bossolasco, S., Pazzi, A., Bestetti, A., Carmillo, P., Wilson, E., McAuliffe, M., et al. (2011). Progressive multifocal leukoencephalopathy (PML) development is associated with mutations in JC virus capsid protein VP1 that change its receptor specificity. J. Infect. Dis. 204, 103–114.

30. Haley, S.A., O’Hara, B.A., Nelson, C.D.S., Brittingham, F.L.P., Henriksen, K.J., Stopa, E.G., and Atwood, W.J. (2015). Human polyomavirus receptor distribution in brain parenchyma contrasts with receptor distribution in kidney and choroid plexus. Am. J. Pathol. 185, 2246–2258.

31. Heiser, K., Nicholas, C., and Garcea, R.L. (2016). Activation of DNA damage repair pathways by murine polyomavirus. Virology 497, 346–356.

32. Inuzuka, T., Ueda, Y., Arasawa, S., Takeda, H., Matsumoto, T., Osaki, Y., Uemoto, S., Seno, H., and Marusawa, H. (2018). Expansion of viral variants associated with immune escape and impaired virion secretion in patients with HBV reactivation after resolved infection. Sci. Rep. 8, 18070.

33. Jelcic, I., Combaluzier, B., Jelcic, I., Faigle, W., Senn, L., Reinhart, B.J., Ströh, L., Nitsch, R.M., Stehle, T., Sospedra, M., et al. (2015). Broadly neutralizing human monoclonal JC polyomavirus VP1-specific antibodies as candidate therapeutics for progressive multifocal leukoencephalopathy. Sci. Transl. Med. 7, 306ra150.

34. Jelcic, I., Jelcic, I., Kempf, C., Largey, F., Planas, R., Schippling, S., Budka, H., Sospedra, M., and Martin, R. (2016). Mechanisms of immune escape in central nervous system infection with neurotropic JC virus variant. Ann. Neurol. 79, 404–418.

35. Jiang, M., Zhao, L., Gamez, M., and Imperiale, M.J. (2012). Roles of ATM and ATR-mediated DNA damage responses during lytic BK polyomavirus infection. PLoS Pathog. 8, e1002898.

36. Justice, J.L., Needham, J.M., and Thompson, S.R. (2019). BK Polyomavirus Activates the DNA Damage Response To Prolong S Phase. J. Virol. 93, e00130–19.

37. Kalinina, T., Iwanski, A., Will, H., and Sterneck, M. (2003). Deficiency in virion secretion and decreased stability of the hepatitis B virus immune escape mutant G145R. Hepatol. Baltim. Md 38, 1274–1281.

38. Kinchen, V.J., Zahid, M.N., Flyak, A.I., Soliman, M.G., Learn, G.H., Wang, S., Davidson, E., Doranz, B.J., Ray, S.C., Cox, A.L., et al. (2018). Broadly Neutralizing Antibody Mediated Clearance of Human Hepatitis C Virus Infection. Cell Host Microbe 24, 717–730.e5.

39. Krautkrämer, E., Klein, T.M., Sommerer, C., Schnitzler, P., and Zeier, M. (2009). Mutations in the BC-loop of the BKV VP1 region do not influence viral load in renal transplant patients. J. Med. Virol. 81, 75– 81.

40. Lauver, M.D., Goetschius, D.J., Netherby-Winslow, C.S., Ayers, K.N., Jin, G., Haas, D.G., Frost, E.L., Cho, S.H., Bator, C., Bywaters, S.M., et al. (2020). Antibody escape by polyomavirus capsid mutation facilitates neurovirulence. ELife 9.

41. Liddington, R.C., Yan, Y., Moulai, J., Sahli, R., Benjamin, T.L., and Harrison, S.C. (1991). Structure of simian virus 40 at 3.8-A resolution. Nature 354, 278–284.

42. Lindner, J.M., Cornacchione, V., Sathe, A., Be, C., Srinivas, H., Riquet, E., Leber, X.-C., Hein, A., Wrobel, M.B., Scharenberg, M., et al. (2019). Human Memory B Cells Harbor Diverse Cross-Neutralizing Antibodies against BK and JC Polyomaviruses. Immunity 50, 668–676.e5.

43. Lukacher, A.E., and Wilson, C.S. (1998). Resistance to polyoma virus-induced tumors correlates with CTL recognition of an immunodominant H-2Dk-restricted epitope in the middle T protein. J. Immunol. Baltim. Md 1950 *160*, 1724–1734.

44. Lynch, R.M., Boritz, E., Coates, E.E., DeZure, A., Madden, P., Costner, P., Enama, M.E., Plummer, S., Holman, L., Hendel, C.S., et al. (2015). Virologic effects of broadly neutralizing antibody VRC01 administration during chronic HIV-1 infection. Sci. Transl. Med. 7, 319ra206.

45. Maginnis, M.S., Ströh, L.J., Gee, G.V., O’Hara, B.A., Derdowski, A., Stehle, T., and Atwood, W.J. (2013). Progressive multifocal leukoencephalopathy-associated mutations in the JC polyomavirus capsid disrupt lactoseries tetrasaccharide c binding. MBio 4, e00247–00213.

46. Maru, S., Jin, G., Desai, D., Amin, S., Shwetank, null, Lauver, M.D., and Lukacher, A.E. (2017). Inhibition of Retrograde Transport Limits Polyomavirus Infection In Vivo. MSphere 2.

47. Mehandru, S., Vcelar, B., Wrin, T., Stiegler, G., Joos, B., Mohri, H., Boden, D., Galovich, J., Tenner-Racz, K., Racz, P., et al. (2007). Adjunctive passive immunotherapy in human immunodeficiency virus type 1-infected individuals treated with antiviral therapy during acute and early infection. J. Virol. 81, 11016–11031.

48. Mockus, T.E., Netherby-Winslow, C.S., Atkins, H.M., Lauver, M.D., Jin, G., Ren, H.M., and Lukacher, A.E. (2020). CD8 T Cells and STAT1 Signaling Are Essential Codeterminants in Protection from Polyomavirus Encephalopathy. J. Virol. 94.

49. Neu, U., Maginnis, M.S., Palma, A.S., Ströh, L.J., Nelson, C.D.S., Feizi, T., Atwood, W.J., and Stehle, T. (2010). Structure-function analysis of the human JC polyomavirus establishes the LSTc pentasaccharide as a functional receptor motif. Cell Host Microbe 8, 309–319.

50. O’Hara, B.A., Gee, G.V., Atwood, W.J., and Haley, S.A. (2018). Susceptibility of Primary Human Choroid Plexus Epithelial Cells and Meningeal Cells to Infection by JC Virus. J. Virol. 92.

51. O’Hara, B.A., Morris-Love, J., Gee, G.V., Haley, S.A., and Atwood, W.J. (2020). JC Virus infected choroid plexus epithelial cells produce extracellular vesicles that infect glial cells independently of the virus attachment receptor. PLoS Pathog. 16, e1008371.

52. Pavlovic, D., Patel, M.A., Patera, A.C., Peterson, I., and Progressive Multifocal Leukoencephalopathy Consortium (2018). T cell deficiencies as a common risk factor for drug associated progressive multifocal leukoencephalopathy. Immunobiology 223, 508–517.

53. Peretti, A., Geoghegan, E.M., Pastrana, D.V., Smola, S., Feld, P., Sauter, M., Lohse, S., Ramesh, M., Lim, E.S., Wang, D., et al. (2018). Characterization of BK Polyomaviruses from Kidney Transplant Recipients Suggests a Role for APOBEC3 in Driving In-Host Virus Evolution. Cell Host Microbe 23, 628–635.e7.

54. Perkins, M.R., Ryschkewitsch, C., Liebner, J.C., Monaco, M.C.G., Himelfarb, D., Ireland, S., Roque, A., Edward, H.L., Jensen, P.N., Remington, G., et al. (2012). Changes in JC virus-specific T cell responses during natalizumab treatment and in natalizumab-associated progressive multifocal leukoencephalopathy. PLoS Pathog. 8, e1003014.

55. Pullen, G.R., Fitzgerald, M.G., and Hosking, C.S. (1986). Antibody avidity determination by ELISA using thiocyanate elution. J. Immunol. Methods 86, 83–87.

56. Qian, M., and Tsai, B. (2010). Lipids and proteins act in opposing manners to regulate polyomavirus infection. J. Virol. 84, 9840–9852.

57. Que, L., Li, Y., Dainichi, T., Kukimoto, I., Nishiyama, T., Nakano, Y., Shima, K., Suzuki, T., Sato, Y., Horike, S., et al. (2021). Interferon-gamma induced APOBEC3B contributes to Merkel cell polyomavirus genome mutagenesis in Merkel cell carcinoma. J. Invest. Dermatol. S0022-202X(21)02636-1.

58. Ray, U., Cinque, P., Gerevini, S., Longo, V., Lazzarin, A., Schippling, S., Martin, R., Buck, C.B., and Pastrana, D.V. (2015). JC polyomavirus mutants escape antibody-mediated neutralization. Sci. Transl. Med. 7, 306ra151.

59. Reid, C.E., Li, H., Sur, G., Carmillo, P., Bushnell, S., Tizard, R., McAuliffe, M., Tonkin, C., Simon, K., Goelz, S., et al. (2011). Sequencing and analysis of JC virus DNA from natalizumab-treated PML patients. J. Infect. Dis. 204, 237–244.

60. Ren, H.M., Kolawole, E.M., Ren, M., Jin, G., Netherby-Winslow, C.S., Wade, Q., Shwetank, null, Rahman, Z.S.M., Evavold, B.D., and Lukacher, A.E. (2020). IL-21 from high-affinity CD4 T cells drives differentiation of brain-resident CD8 T cells during persistent viral infection. Sci. Immunol. 5, eabb5590.

61. Sanjuán, R., and Domingo-Calap, P. (2016). Mechanisms of viral mutation. Cell. Mol. Life Sci. 73, 4433–4448.

62. Schneider-Hohendorf, T., Rossaint, J., Mohan, H., Böning, D., Breuer, J., Kuhlmann, T., Gross, C.C., Flanagan, K., Sorokin, L., Vestweber, D., et al. (2014). VLA-4 blockade promotes differential routes into human CNS involving PSGL-1 rolling of T cells and MCAM-adhesion of TH17 cells. J. Exp. Med. 211, 1833–1846.

63. Son, Y.M., Cheon, I.S., Wu, Y., Li, C., Wang, Z., Gao, X., Chen, Y., Takahashi, Y., Fu, Y.-X., Dent, A.L., et al. (2021). Tissue-resident CD4+ T helper cells assist the development of protective respiratory B and CD8+ T cell memory responses. Sci. Immunol. 6, eabb6852.

64. Sowd, G.A., Li, N.Y., and Fanning, E. (2013). ATM and ATR activities maintain replication fork integrity during SV40 chromatin replication. PLoS Pathog. 9, e1003283.

65. Stehle, T., and Harrison, S.C. (1997). High-resolution structure of a polyomavirus VP1-oligosaccharide complex: implications for assembly and receptor binding. EMBO J. 16, 5139–5148.

66. Stüve, O., Marra, C.M., Bar-Or, A., Niino, M., Cravens, P.D., Cepok, S., Frohman, E.M., Phillips, J.T., Arendt, G., Jerome, K.R., et al. (2006). Altered CD4+/CD8+ T-cell ratios in cerebrospinal fluid of natalizumab-treated patients with multiple sclerosis. Arch. Neurol. 63, 1383–1387.

67. Sunyaev, S.R., Lugovskoy, A., Simon, K., and Gorelik, L. (2009). Adaptive mutations in the JC virus protein capsid are associated with progressive multifocal leukoencephalopathy (PML). PLoS Genet. 5, e1000368.

68. Swimm, A.I., Bornmann, W., Jiang, M., Imperiale, M.J., Lukacher, A.E., and Kalman, D. (2010). Abl family tyrosine kinases regulate sialylated ganglioside receptors for polyomavirus. J. Virol. 84, 4243– 4251.

69. Tokonami, N., Takata, T., Beyeler, J., Ehrbar, I., Yoshifuji, A., Christensen, E.I., Loffing, J., Devuyst, O., and Olinger, E.G. (2018). Uromodulin is expressed in the distal convoluted tubule, where it is critical for regulation of the sodium chloride cotransporter NCC. Kidney Int. 94, 701–715.

70. Toma, J., Weinheimer, S.P., Stawiski, E., Whitcomb, J.M., Lewis, S.T., Petropoulos, C.J., and Huang, W. (2011). Loss of asparagine-linked glycosylation sites in variable region 5 of human immunodeficiency virus type 1 envelope is associated with resistance to CD4 antibody ibalizumab. J. Virol. 85, 3872–3880.

71. Tremolada, S., Delbue, S., Castagnoli, L., Allegrini, S., Miglio, U., Boldorini, R., Elia, F., Gordon, J., and Ferrante, P. (2010). Mutations in the external loops of BK virus VP1 and urine viral load in renal transplant recipients. J. Cell. Physiol. 222, 195–199.

72. Trkola, A., Kuster, H., Rusert, P., Joos, B., Fischer, M., Leemann, C., Manrique, A., Huber, M., Rehr, M., Oxenius, A., et al. (2005). Delay of HIV-1 rebound after cessation of antiretroviral therapy through passive transfer of human neutralizing antibodies. Nat. Med. 11, 615–622.

73. Van Loy, T., Thys, K., Ryschkewitsch, C., Lagatie, O., Monaco, M.C., Major, E.O., Tritsmans, L., and Stuyver, L.J. (2015). JC virus quasispecies analysis reveals a complex viral population underlying progressive multifocal leukoencephalopathy and supports viral dissemination via the hematogenous route. J. Virol. 89, 1340–1347.

74. Vaz, B., Cinque, P., Pickhardt, M., and Weber, T. (2000). Analysis of the transcriptional control region in progressive multifocal leukoencephalopathy. J. Neurovirol. 6, 398–409.

75. Wilson, J.J., Lin, E., Pack, C.D., Frost, E.L., Hadley, A., Swimm, A.I., Wang, J., Dong, Y., Breeden, C.P., Kalman, D., et al. (2011). Gamma interferon controls mouse polyomavirus infection in vivo. J. Virol. 85, 10126–10134.

76. Wilson, J.J., Pack, C.D., Lin, E., Frost, E.L., Albrecht, J.A., Hadley, A., Hofstetter, A.R., Tevethia, S.S., Schell, T.D., and Lukacher, A.E. (2012). CD8 T cells recruited early in mouse polyomavirus infection undergo exhaustion. J. Immunol. Baltim. Md 1950 *188*, 4340–4348.

77. Wollebo, H.S., White, M.K., Gordon, J., Berger, J.R., and Khalili, K. (2015). Persistence and pathogenesis of the neurotropic polyomavirus JC. Ann. Neurol. 77, 560–570.

78. Zhao, L., and Imperiale, M.J. (2021). A Cell Culture Model of BK Polyomavirus Persistence, Genome Recombination, and Reactivation. MBio e0235621.

79. Zheng, H.-Y., Takasaka, T., Noda, K., Kanazawa, A., Mori, H., Kabuki, T., Joh, K., Oh-Ishi, T., Ikegaya, H., Nagashima, K., et al. (2005a). New sequence polymorphisms in the outer loops of the JC polyomavirus major capsid protein (VP1) possibly associated with progressive multifocal leukoencephalopathy. J. Gen. Virol. 86, 2035–2045.

80. Zheng, H.-Y., Ikegaya, H., Takasaka, T., Matsushima-Ohno, T., Sakurai, M., Kanazawa, I., Kishida, S., Nagashima, K., Kitamura, T., and Yogo, Y. (2005b). Characterization of the VP1 loop mutations widespread among JC polyomavirus isolates associated with progressive multifocal leukoencephalopathy. Biochem. Biophys. Res. Commun. 333, 996–1002.

