## Supplemental Tables/Figs for "T Cell Deficiency Precipitates Antibody Evasion and Emergence of Neurovirulent Polyomavirus"

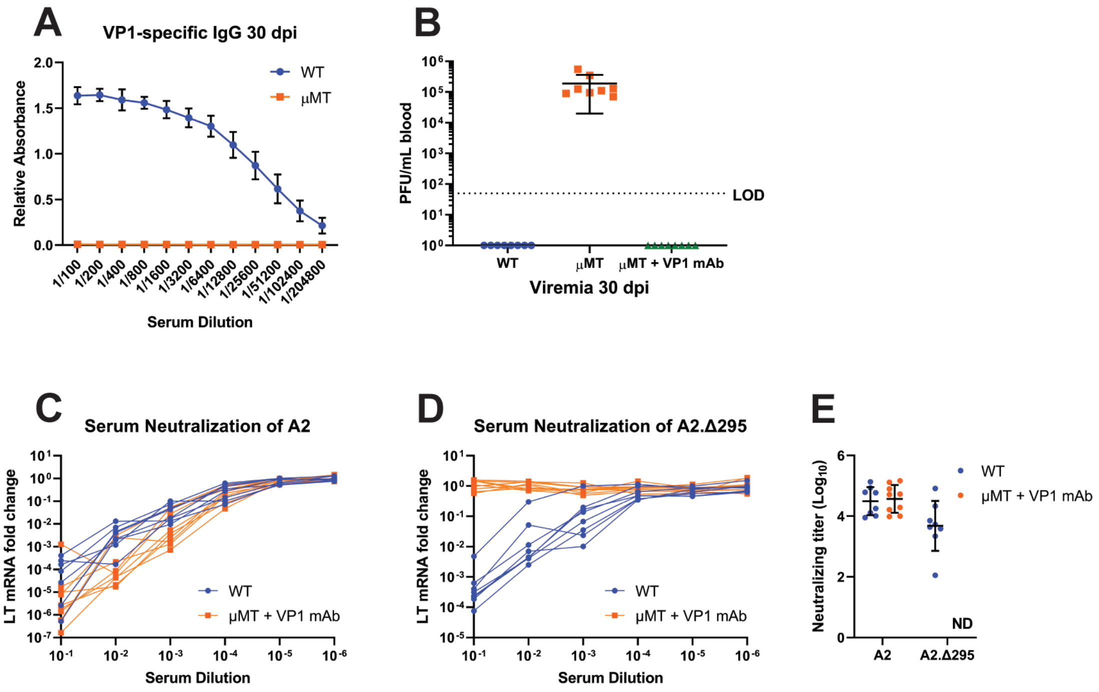
A
VP1-specific IgG 30 dpi
Relative Absorbance
WT
μMT
1/100 1/200 1/400 1/800 1/1600 1/3200 1/6400 1/12800 1/25600 1/51200 1/102400 1/204800
Serum Dilution
B
PFU/mL blood
LOD
WT
μMT
μMT + VP1 mAb
Viremia 30 dpi
C
Serum Neutralization of A2
LT mRNA fold change
WT
μMT + VP1 mAb
Serum Dilution
D
Serum Neutralization of A2.Δ295
LT mRNA fold change
WT
μMT + VP1 mAb
Serum Dilution
E
Neutralizing titer (Log10)
WT
μMT + VP1 mAb
ND
A2
A2.Δ295

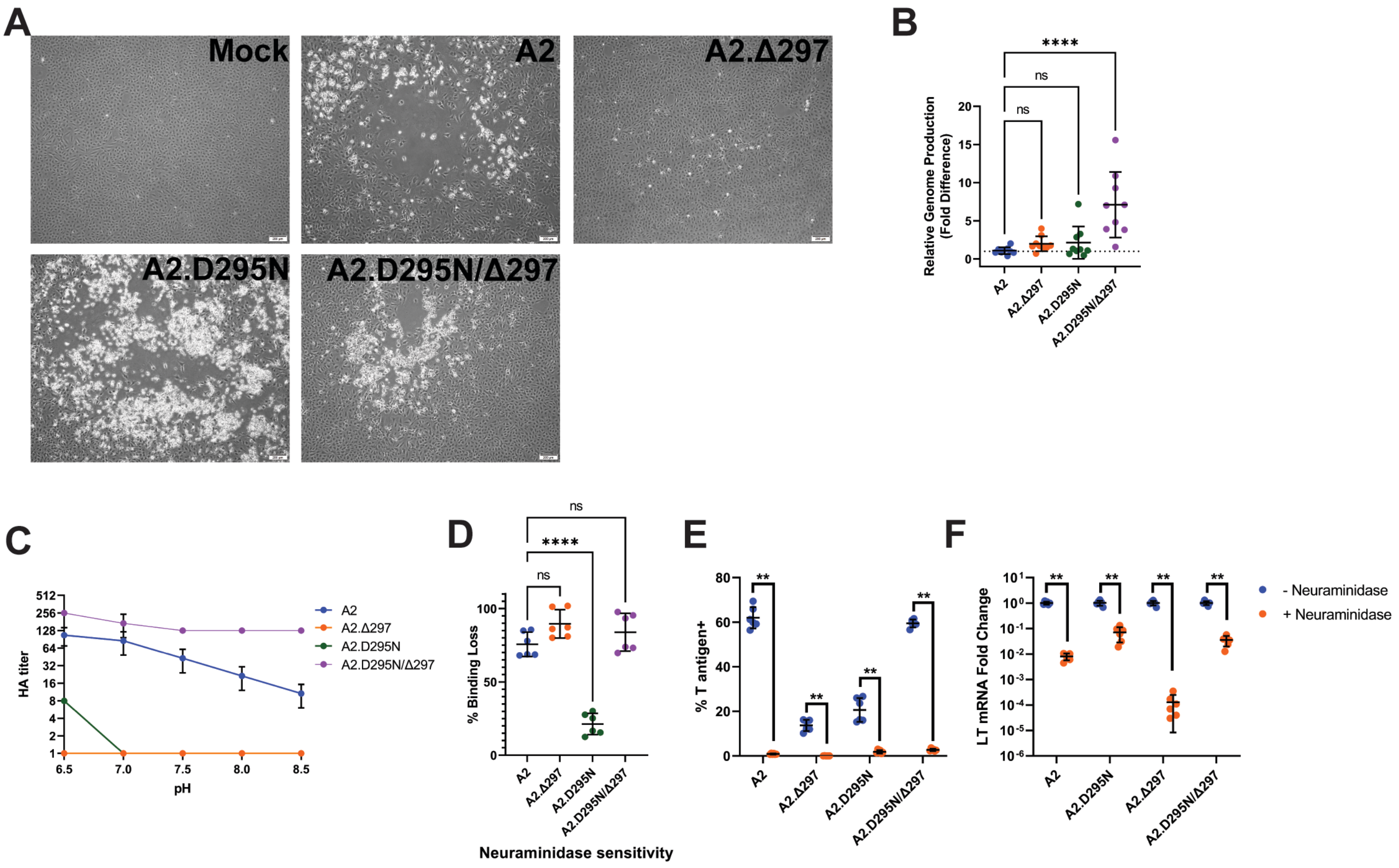
A
Mock
A2
A2.Δ297
A2.D295N
A2.D295N/Δ297
B
Relative Genome Production (Fold Difference)
****
ns
ns
A2
A2.Δ297
A2.D295N
A2.D295N/Δ297
C
HA titer
pH
A2
A2.Δ297
A2.D295N
A2.D295N/Δ297
D
% Binding Loss
ns
****
ns
Neuraminidase sensitivity
E
% T antigen+
**
F
LT mRNA Fold Change
**
- Neuraminidase
+ Neuraminidase

**Table S1**. **Identity, location, and frequency of detected VP1 mutations, related to Figure 3.**

| Condition | VP1 Mutation | Loop | # of mice |
|---|---|---|---|
| IgG | E68KΔ297 | BC,HI | 1 |
| | E91VΔ297 | BC,HI | 1 |
| | Δ145-150Δ294 | DE,HI | 1 |
| | E187KΔ297 | EF, HI | 1 |
| | Δ294 | HI | 1 |
| | Δ295 | HI | 3 |
| | V296F | HI | 1 |
| αCD4 + αCD8β | Δ292Y294H | HI | 1 |
| | Y294D | HI | 1 |
| | Δ295 | HI | 7 |
| | D295AΔ297 | HI | 1 |
| | D295NΔ297 | HI | 1 |
| αCD4 | E68KΔ297 | BC,HI | 1 |
| | E87KΔ297 | BC,HI | 1 |
| | N293KΔ297 | HI | 2 |
| | Δ295 | HI | 2 |
| | D295NΔ297 | HI | 1 |
| αCD8β | Δ295 | HI | 2 |

**Table S2. Oligonucleotide Sequences.**

| Designation | Reference | Sequence 5’-3’ |
|---|---|---|
| Δ292/Y294H Forward | This paper | GGGAAGCCCTCTCCAGTGATGGACATCATGGTTTGT AACTCTCCAGCCCATTATATC |
| Δ292/Y294H Reverse | This paper | GATATAATGGGCTGGAGAGTTACAAACCATGATGTC CATCACTGGAGAGGGCTTCCC |
| Y294D Forward | This paper | GGGAAGCCCTCTCCAGTGATGGACATCATCGTTTCT TGTAACTCTCCAGCCC |
| Y294D Reverse | This paper | GGGCTGGAGAGTTACAAGAAACGATGATGTCCATC ACTGGAGAGGGCTTCCC |
| D295A/Δ297 Forward | This paper | GGGAAGCCCTCTCCAGTGGACAGCATAGTTTCTTGT AACTCTCCAGCCCATTATATC |
| D295A/Δ297 Reverse | This paper | GATATAATGGGCTGGAGAGTTACAAGAAACTATGCT GTCCACTGGAGAGGGCTTCCC |
| D295N/Δ297 Forward | This paper | CTGGGAAGCCCTCTCCAGTGGACATTATAGTTTCTT GTAACTCTCCAGCCC |
| D295N/Δ297 Reverse | This paper | GGGCTGGAGAGTTACAAGAAACTATAATGTCCACTG GAGAGGGCTTCCCAG |
| D295N Forward | This paper | GGGAAGCCCTCTCCAGTGATGGACATTATAGTTTCT TGTAACTCTCCAGCCC |
| D295N Reverse | This paper | GGGCTGGAGAGTTACAAGAAACTATAATGTCCATCA CTGGAGAGGGCTTCCC |
| Δ297 Forward | This paper | GGGAAGCCCTCTCCAGTGGACATCATAGTTTCTTGT AACTCTCCAGCCCATTATATC |
| Δ297 Reverse | This paper | GATATAATGGGCTGGAGAGTTACAAGAAACTATGA TGTCCACTGGAGAGGGCTTCCC |
| VP1 TA cloning Forward | This paper | GGTCAACATAGCGCGTCATA |
| VP1 TA cloning Reverse | This paper | CCAGTTGAAATCTGGCATCC |
| VP1 sequencing | This paper | TACACTCTAACCTCCTCTACCTG |
| LT DNA qPCR Forward | Wilson et al., 2012 | CGCACATACTGCTGGAAGAAGA |
| LT DNA qPCR Reverse | Wilson et al., 2012 | TCTTGGTCGCTTTCTGGATACAG |
| LT DNA qPCR probe | Wilson et al., 2012 | ATCCTTGTGTTGCTGAGCCCGATG |
| LT mRNA qPCR Forward | Maru et al., 2017 | AGGAATTGAACAGTCTCTGGG |
| LT mRNA qPCR Reverse | Maru et al., 2017 | GTCATCGTGTAGTGGACTGTG |
| LT mRNA qPCR probe | Maru et al., 2017 | AACCGGCTTCCAGGGCTCT |
| VP1 Amplification Forward | Lauver et al., 2020 | CGACCCCTTGAAGGACATATGTGAA |
| VP1 Amplification Reverse | Lauver et al., 2020 | CACCTACTTGGGCAACAGTCA |
